# Seasonal patterns in b-vitamins and cobalamin co-limitation in the Northwest Atlantic

**DOI:** 10.1101/2024.11.10.622835

**Authors:** Catherine Bannon, Patrick L. White, Elden Rowland, Kiran J. More, Anna Gleason, Megan Roberts, Emmanuel Devred, Lindsay Beazley, Julie LaRoche, Erin M. Bertrand

## Abstract

B-vitamins are important co-enzymes that have long been hypothesized to play key roles in marine ecosystems. However, environmental measurements remain scarce, which limits our understanding of their potential impact. Here, we present mass spectrometry-based measurements of b-vitamins (B_1_, B_2_, B_3_, B_5_, B_6,_ B_12_) and related vitamers along a transect in the Northwest Atlantic Ocean, in both particulate phase and dissolved in seawater, seasonally over 5 years, and couple this with targeted investigations of the impact of B_12_ (cobalamin) on phytoplankton growth. We show that these metabolites are present at femto to pico-molar concentrations and demonstrate that season explains most variance in particulate phase b-vitamins but not dissolved, offering further evidence that metabolite inventories in these two phases are often decoupled. We find correlations between particulate organic carbon with particulate B_1_ and B_3_, and between chlorophyll *a* and particulate B_2_ and DMB in fall but not spring, indicating unique seasonal drivers of vitamin inventories. Of all measured vitamins, only cobalamin was enriched in the particulate over dissolved phase, predominantly in spring. We documented nitrogen and cobalamin co-limitation of phytoplankton growth during spring bloom decline, when dissolved cobalamin is seemingly drawn down, but not during fall, when dissolved cobalamin concentrations remain elevated. These seasonal differences may be underpinned by the increased importance of cobalamin remodeling and recycling during the fall. This study provides insights into the absolute concentrations, stoichiometry, and variability of b-vitamins in the ocean and offers evidence that cobalamin exerts seasonally-varying controls on Northwest Atlantic marine ecosystems.

## Introduction

B-vitamins like cobalamin (B_12_), thiamine (B_1_), biotin (B_7_), riboflavin (B_2_), niacinamide/niacin (B_3_), pantothenic acid (B_5_), pyridoxine (B_6_) and folate (B_9_), are facilitator metabolites (Moran et al. 2022) required as co-factors in central metabolism. In the ocean, b-vitamins have been hypothesized, for over a century, to have the power to influence microbial community composition and activity (Allen 1914; Droop 1954; Carlucci and Bowes 1970). Their suggested impact, in part, is attributed to b-vitamin auxotrophy, or the absolute requirement for an exogenous source of the b-vitamin, which is expected to be widespread in marine microbial communities (Croft et al. 2006; Gómez-Consarnau et al. 2018; Cooper et al. 2019; Gregor et al. 2023, Bayer et al, 2024). Such evidence points towards b-vitamins being key microbial currencies with a dynamic inventory in the ocean. However, their roles and impacts in marine ecosystems remains unclear, in part due to complications associated with measuring them.

Obtaining reliable measurements of b-vitamins and vitamers (e.g., precursors, degradation products, related compounds) is difficult due to matrix effects, low concentrations, and chemical diversity (Kido Soule et al. 2015; Moran et al. 2022). There are just a few examples of measurements of b-vitamins (i.e. only B_2_ and B_5_) or one b-vitamin and a suite of related compounds (i.e. thiamine and related compounds) over space (Heal et al. 2017; Suffridge et al. 2020; Johnson et al. 2023) and time (year: (Longnecker et al. 2024), day: (Boysen et al. 2021), season: (Bittner et al. 2024)). Reliably measuring a suite of b-vitamins over both space and time could provide further insights into the apparent interconnectedness of b-vitamins on a cellular (Bertrand and Allen 2012; Monteverde et al. 2017) and community scale (Cooper et al. 2019). Additionally, most measurements to date have been made in either particulate (cell, endo-metabolome) or dissolved (seawater, exo-metabolome) phase, despite evidence that these pools do not always correlate, and that unique information is obtained from each pool (Johnson et al. 2023). Dissolved metabolites, which require time-consuming solid phase extraction steps to concentrate metabolites before analysis, represent the pool of public goods available to the microbial community, and are influenced by both biotic (e.g. uptake, release) and abiotic (e.g. light, temperature, hydrolysis) factors. Particulate measurements provide a snapshot of that microbial community’s intracellular contents which may reflect composition and activity. Investigating b-vitamins in both phases, over time and space in a well-characterized ocean region could provide new insights into environmental b-vitamin dynamics and the ways in which these vitamins may impact ecosystem processes.

Elucidating the environmental dynamics of cobalamin, vitamin B_12_, is of particular interest because of its ability to co-limit primary production and influence community composition in various parts of the ocean (Bannon et al. 2022). Cobalamin is a cobalt-containing corrin ring with DMB (5,6-dimethylbenzimidazole) at its lower axial ligand and various upper axial ligands (Me, Ado, OH, CN-B_12_). Me and Ado-B_12_ are the biologically active co-factors (Banerjee and Ragsdale 2003), while CN-B_12_ is a synthetic form that is not known to be biologically produced (Martens et al. 2002; Warren et al. 2002). These forms are photolabile and convert into OH-B_12_ on the scale of seconds to days, and as such, OH-B_12_ is generally expected to be the dominant dissolved form in sunlit seawater (Bannon et al. 2024). Only select bacteria are able to produce cobalamin *de novo*, however, key eukaryotic phytoplankton are cobalamin auxotrophs (Bertrand et al. 2013) and non-auxotrophic facultative consumers are also significant cobalamin sinks in the ocean (Gleason et al, in prep). Most cyanobacteria produce and use their own cobalamin analog, pseudocobalamin (psB_12_), which can be remodeled into cobalamin by a subset of organisms (Helliwell et al, 2016). Cobalamin remodeling remains poorly studied *in situ*, limiting our ability to determine its impact on cobalamin availability. Understanding patterns in cobalamin availability, use, limitation, and remodeling is of great interest for efforts to identify a theoretical framework that predicts how this micronutrient impacts ecosystems.

The Scotian Shelf and Slope (SSS) situated off Nova Scotia, eastern Canada, is an oceanographically complex region influenced by both cold fresh water from the Nova Scotia Current (originated in the Gulf of St. Lawrence) and Shelf Break Current (extension of the Labrador Current), and the flow of warm salty water from the Gulf Stream (Hannah et al, 2000). The shelf is a wide (up to 240 km) but shallow (average ∼100 m) bank embedded with deep basins and channels that breaks to the slope where depth drops to >3000 m. Seasonal, interannual, and decadal variability in the physical, chemical, and lower trophic-level biological characteristics of the SSS has been monitored twice annually by Fisheries and Oceans Canada’s Atlantic Zone Monitoring Program since 1998 (AZMP; Therriault et al. 1998). Such work has demonstrated that the SSS follows the canonical North Atlantic annual cycle, where a diatom-dominated bloom is induced in spring followed by an oligotrophic, cyanobacteria-dominated fall bloom (Li et al. 2006), supporting distinct seasonal microbial communities (Zorz et al. 2019; Robicheau et al. 2022). To date, measurements of b-vitamins and vitamers in this region are limited to that of thiamine-related compounds in the dissolved phase (Paerl et al. 2023) and particulate cobalamins (Bannon et al. 2024) from a limited number of stations and years. Previous work leveraged metagenomic analyses to identify potential prokaryotic cobalamin producers, remodelers, and consumers in both spring and fall (Soto et al. 2023). This work, along with the long-term oceanographic monitoring conducted by the AZMP, makes the SSS an ideal region to study variability of b-vitamins *in situ* and further elucidate cobalamin dynamics.

Here, we perform quantitative measurements of b-vitamins (B_1_, B_2_, B_3_, B_5_, B_6,_ B_12_) and vitamers (DMB, HET - 4-methyl-5-thiazoleethanol, HMP - 4-amino-5- hydroxymethyl-2-methylpyrimidine, FAMP - N-formyl-4-amino-5-aminomethyl-2- methylpyrimidine) in both particulate and dissolved samples at different depths (1 to 250 m) from a fixed CTD transect positioned across the SSS in the North Atlantic Ocean (NWA), sampled during the spring and fall over 5 years from 2016 to 2020. We then examine trends between and within the particulate and dissolved phases to investigate the variability of b-vitamins and begin to uncover factors that influence them. Finally, we examine the role of cobalamins in the region by combining these measurements with nutrient addition bioassays to examine seasonal patterns in cobalamin dynamics.

## Materials and methods

### Shipboard sampling

We collected environmental samples from 14 fixed stations on the AZMP’s Halifax Line (HL01, HL02, HL03, HL03.3, HL04, HL05.5, HL06, HL06.3, HL07, HL08, HL09, HL10, HL11, HL12) during 8 AZMP monitoring surveys over 5 years (2016 – 2020). The surveys were conducted primarily from the Canadian Coast Guard oceanographic research vessel *Hudson* during which discrete water samples were collected at a subset of standard depths (near-surface, 20, 40, 50, 60, 80, 100, 250) via the deployment of a SeaBird Electronics 911 CTD-Rosette system outfitted with 10 – 12 L Niskin bottles. For vitamin analysis, we collected ∼1 L of water from 4 depths at each station, where depths chosen varied based on bathymetry. Samples were gently filtered through a 0.2 μm Nylon filter into an amber glass bottle in the dark to minimize photodegradation. Two 500 mL filtrate samples were collected in acid-washed, milli-Q water rinsed, and sample-rinsed amber high-density polyethylene (HDPE) bottles then frozen at -20 °C until processing for dissolved metabolite analysis. A total of 148 particulate metabolite samples and duplicate extractions of 133 dissolved samples were processed, as described below.

CTD salinity and temperature (°C), sample time, nutrient concentration (phosphate, nitrate, ammonia) from bottle measurements (mmol m^-3^), chlorophyll *a (*chl *a*) concentration (fluorometry following Holm-Hansen method) (mg m^-3^), and quantitative pigment data (mg m^-3^) (High Performance Liquid Chromatography, HPLC) from the AZMP cruises were provided by the Bedford Institute of Oceanography (BIO).

Measurements of particulate organic carbon (POC, mg m^-3^), particulate organic nitrogen (PON, mg m^-3^) and HPLC pigments (mg m^-3^) were restricted to 1 m depth. Data were collected and quality-controlled as described elsewhere (Mitchell et al. 2002; Devine et al. 2014).

### Extractions

Due to the extreme photolability of cobalamins (Bannon et al. 2024), we performed all extractions (particulate and dissolved) in a dark, windowless room, with a red headlamp as the only light source.

#### Particulate extractions

Particulate metabolite extraction was performed following the methodology described by Heal et al. (2017) with minor modifications. Briefly, particulate samples were spiked with 1 picomol of heavy-CN-B_12_ (Cambridge Isotopes Labs, CLM-9770-E) internal standard then extracted using an organic solvent extraction (40:20:20 acetonitrile: methanol: MQ water) paired with bead beating. Solvent was removed under vacuum (Eppendorf, Mississauga, ON) at room temperature and samples were resuspended with 100 µL of buffer A (H_2_O containing 0.1% formic acid). All samples were vortexed and centrifuged at 10,000 x g for 3 min at 4 °C and then diluted with buffer A in conical polypropylene HPLC vials (Phenomenex, Torrance, CA) by 3-fold for analysis on the C18 Acclaim prior to mass spectrometry analysis.

#### Dissolved extractions

One dissolved sample (20 mL) was extracted from each duplicate filtrate samples on a 500 mg HyperSep C18 solid-phase extraction (SPE) cartridges (ThermoScientific) using a vacuum manifold. SPE cartridges were preconditioned with 2 x 2 mL MeOH then 2 x 2 mL deionized water and kept wet through the entire protocol.

Samples were extracted at ∼1 mL per min, washed with 2 x 2 mL milli-Q water then eluted with 2 x 0.85 mL MeOH into microtubes. Eluent was dried for ∼2 h under vacuum (Vacufuge, Eppendorf), in the dark, then stored at -80 °C prior to analysis. Each sample was resuspended with 100 µL of buffer A before mass spectrometry analysis. Percent recovery was determined by spiking 20 mL seawater samples with each analyte before or after preconcentration on SPE (n = 9). Only metabolites with an average percent recovery of >40% were included in this study. These are typical percent recoveries in seawater due to the complexity of the samples (Johnson et al. 2017). All reported concentrations and limits of detection (pM) are corrected for percent recoveries.

### Mass spectrometry analysis

All particulate and dissolved samples were analyzed on a Dionex Ultimate-3000 LC system coupled to the electrospray ionization source of a TSQ Quantiva triple-stage quadrupole mass spectrometer (ThermoFisher) operated in selected reaction monitoring mode, with the following settings: Q1 and Q3 resolution 0.7 (FWHM), 6 ms dwell time, CID Gas 2.5 mTorr, spray voltage in 3500 V positive ion mode, sheath gas 6, auxiliary gas 2, ion transfer tube temperature 325 °C, vaporizer temperature 100 °C. Duplicate 5 μL injections were performed onto an Acclaim PepMapTM RSLC, 300 μm x 150 mm, C18, 2 μm, 100 Å (PN164537, ThermoScientific) column held at 50 °C and subject to a gradient of 4–99% B over 8 minutes (A: 0.1% formic acid in H2O; B: 0.1% formic acid in MeCN) at 10 μL per min. The total run time including washing and equilibration was 12 min using transition list reported in Table S1.

### Approach to quantification

Matrix effects of environmental metabolomic samples are known to be severe (Sysi-Aho et al. 2007; Kido Soule et al. 2015; Boysen et al. 2021). As such, various approaches were taken in this study to minimize the impact of matrix effects on analyte quantification.

Samples were grouped based on previously reported similarity of microbial communities (Craig et al. 2015; Zorz et al. 2019; Robicheau et al. 2022) and physicochemical characteristics (Fratantoni and Pickart 2007; Shadwick and Thomas 2014). Particulate and dissolved samples were separated into four matrix-specific groups, fall and spring, on and off-shelf (Fig S1). We combined equal portions of each sample to obtain a quality control (QC) pool of each matrix group that was then used as an intra-run QC injection and the base for calibration curves, adjusted for fold-dilution. Matrix-group specific calibration curves were performed in duplicate injections of 2.5, 5, 25, 125 fmol on analytical column for all forms of B_12_, DMB, HET and B_2_; 12.5, 25, 250 fmol on analytical column for B_1_, B_3_ (niacin and niacinamide), HMP, FAMP, B_5_, B_6_.

Details of metabolite standards are provided in the supplemental methods. Calibrations curves were used for quantification if R^2^ were all >74%. Limits of detection (LOD) were determined according to Armbruster and Pry (2008) based on the fmol analyte on analytical column. Slopes and LODs calculated with each matrix-group specific calibration curve were then applied to the respective matrix group (Fig S1).

Heavy-CN-B_12_ spikes onto each particulate filter prior to extraction has been previously shown to minimize the impact of matrix effects and account for variability introduced throughout analysis (Heal et a, 2014). During quantification, all B_12_ forms were normalized to the heavy-CN-B_12_ signal for analysis (Fig S2) to account for human errors during extraction and instrument variability.

Finally, samples were further visually inspected and analyzed following quality control criteria modified from Boysen et al. 2018. Metabolite concentrations were reported in samples if, (i) the peak had the same retention time (± 0.2 min) as the authentic standard, (ii) at least two daughter fragments were present with co-occurring peak, and (iii) daughter fragments were present in the same order of intensity as authentic standard. Raw files generated with Xcalibur software (ThermoFisher) were uploaded into Skyline-daily (version 23.1.1.459, University of Washington) and the transitions with the best signal-to-noise and lowest interference were selected for quantification reported in Table S1.

### Co-limitation nutrient addition experiments

Water samples from 5 m depth were collected on October 5^th^, 2016 (HUD2016027) and May 3^rd^, 2017 (COR2017001) from HL02 via CTD sampling rosette. Water samples were transferred into carboys, stored in a cooler and transported to the laboratory where acid cleaned 300 mL polycarbonate bottles were used to set up an incubation experiment in triplicate (spring) and quadruplicate (fall) for control, +N, +B_12_, and +N+B_12_ treatments within 24 hours of collection. Final concentrations of nitrate (NaNO_3_, Sigma-Aldrich) and B_12_ (CN-B_12_, Fisher Bioreagents) added to the bottles were 10 μmol L^-1^ and 100 pmol L^-1^ respectively. The bottles were then incubated at 4.5 °C (spring incubation) and 18 °C (fall incubation) under 90-110 µE/m^2^/second light 12:12 light dark cycle. In vivo chl *a* concentration measurements, a proxy for phytoplankton growth, were taken every 24 hours, using a Turner Designs AquaFluor Handheld Fluorometer.

For the spring incubation, flow cytometry was performed daily to obtain cell counts of photosynthetic eukaryotes (size groups: <3 µm, >3 µm <10 µm, and >10 µm), cryptophytes, non-photosynthetic bacteria, and *Synechococcus* as described elsewhere (Robicheau et al. 2022). Briefly, 2 mL of each replicate was collected and preserved in 1% PFA for 10 minutes before placing into liquid N_2_ and stored at -20 °C until analysis. For analysis, samples were prefiltered (35 μm) then stained with SYBR-Green prior to recording autofluorescent cell counts on a CSampler-equipped BD AccuriTM C6 Flow Cytometer (BD Biosciences, USA) with optical filters for chlorophyll [>670 nm] and phycoerythrin [585/40 nm] detection. Cell counts were corrected using blanks (0.2 μm filtered seawater).

After four (spring) and five (fall) day incubation periods, 100 mL volumes from each bottle were filtered (Whatman GF/F) and stored at -80 °C until processed for chl *a* extraction. Extracted chl *a* quantification was performed using EPA method 445 with a Turner Designs AquaFluor Handheld Fluorometer and Turbidimeter (Arar and Collins 1997). Prior to taking measurements, calibration was done by creating a calibration curve using chl *a* (Sigma from spinach) in dilutions of 0.2, 2, 5, 20, and 200 μg chl *a* L^-1^.

### Statistical analyses

A quantitative multivariate regression tree (MRT) analysis was performed to investigate which variables accounted for the variance in b-vitamin datasets using “mvpart” r package (De’ath 2002). MRT is robust to missing (or zero) values, does not assume a linear relationship, handles raw, unnormalized data, as well as continuous and categorical data. B-vitamins and vitamers data were scaled around the mean prior to MRT analysis and only b-vitamins and vitamers that were reliably detected in all matrix groups were used. Inputs to the analysis were the quantitative b-vitamin and vitamer measurements and corresponding values for season, location, depth, salinity, temperature, nitrate, phosphate, ammonia, time of day, and year.

Hierarchical clustering was performed to investigate the similarity across samples and metabolites in both particulate and dissolved phase. For this analysis, b-vitamins and vitamers that fell bellow LOD were set to zero. Data were scaled around the mean prior to analysis and Euclidean distances were calculated with r package “vegan”. All dendrograms were generated using stats package and the “ward-D2” method.

A Spearman correlation matrix was used to identify linear and monotonic relationships in R using the package “corrplot”. As this analysis is sensitive to zero values, b-vitamin concentrations that fell below LOD were set to the average LOD across all matrix-specific groups. Relationships are shown if significance level was less than 0.01. Specific relationships were further interrogated for seasonal differences using linear models.

Data acquired from bottle incubation assay (chl *a* quantification and flow cytometry counts) were assessed for normality (Shapiro-Wilk test), homogeneous in variance (Levene test), and determined significant using an Analysis of Variance (ANOVA). Tukey HSD was used for post hoc test and significance was determined if p- value < 0.05. For bottle incubation assays, paired t-tests were also run post hoc and determined significant if p-value < 0.05.

## Results

### Oceanographic context

Water temperature was warmer in fall (average: 13.6 °C) compared to spring (average: 6.8 °C) both on and off shelf (Fig 1B). Salinity was similar in both seasons but was higher off-shelf (average: 34.5) compared to on-shelf (average: 32.0) due to the proximity to the Gulf Stream (Fig 1C). In general, nitrate concentrations were higher at 20-80 m than 1-20 m in both seasons and locations. In the surface water, nitrate was generally lower in fall (average: 0.3 mmol m^-3^) than in spring (average: 2.0 mmol m^-3^) particularly in off-shelf samples (Fig 1D). In spring, chl *a* concentrations were in average 1.12 mg m^-3^ and 0.37 mg m^-3^ in fall (Fig 1E), consistent with the recurring annual spring bloom that varies in intensity throughout the time series and is followed by small increases in chl *a* in the fall (Fig 1F). Spring sampling often coincided with the decline of the bloom, with the exception of the spring cruise in 2018 and 2019 where the peak of the bloom was sampled.

**Figure 1:**
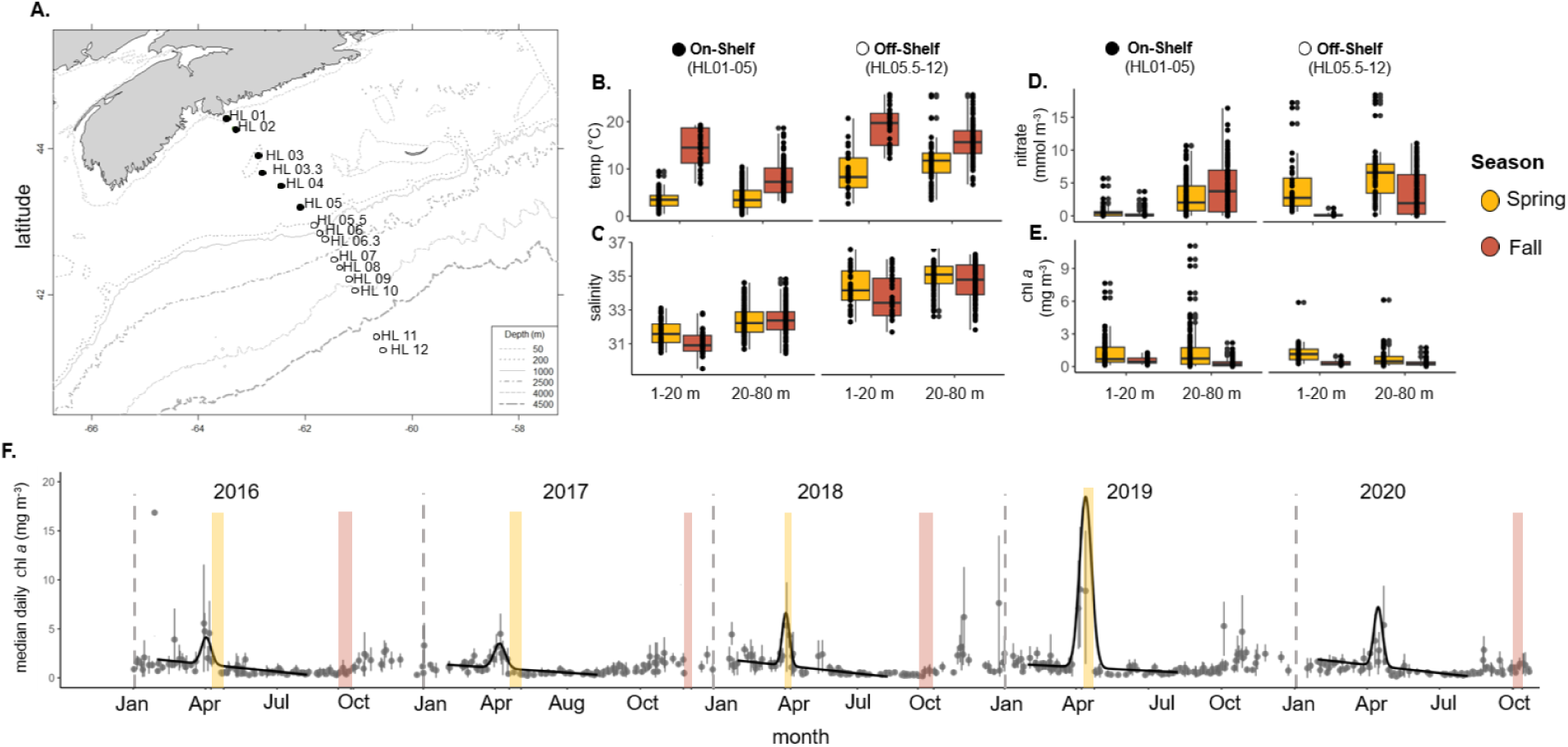
Northwest Atlantic oceanographic context from 2016-2020. (A) Map of Halifax Line stations (HL01-HL12) in the NWA sampled over 2016-2020. Boxplots of (B) temperature (°C), (C) salinity, (D) nitrate (mmol m^-3^), and (E) chl *a* (mg m^-3^) at discrete depths where vitamin samples were obtained, from 1-20 m and 20-80 m depth on (HL01-05) and off (HL05.5-12) shelf during spring (yellow) and fall (orange). (F) Satellite-derived median daily chl *a* concentration from 2016-2020 over time at HL02 using moderate resolution imaging spectroradiometer (MODIS) data (Clay et al. 2021). Black line indicates the shifted gaussian model fits, and timing of the eight research cruises included in this study are highlighted in yellow (spring) and orange (fall).

### Detection, quantification and trends of particulate and dissolved b-vitamins and vitamers on the Scotian Shelf and Slope

#### Particulate samples

Concentrations of 11 vitamin and vitamer metabolites (Me-B_12_, Ado-B_12_, OH-B_12_, CN-B_12_, DMB, B_1_, HET, HMP, B_2_, B_3_, B_5_) were quantified in this study (Table 1, Fig 2A). Limit of detections (LOD) were highest for B_1_, B_3_, and B_5_ but ranged within the femtomolar range for all cobalamin forms and DMB. Quantification of HMP was restricted to select matrix groups as some of the calibration curves failed to pass quality control (>74% R^2^). Significant matrix effects were accounted for by performing matrix group specific quantification in particulate samples. Calibration curves, and resultant slopes, were particularly variable between matrix groupings for early eluting metabolites like B_3_ (slope = 2,100 ± 1,400) compared to vitamers with later retention times like DMB (slope = 40,000 ± 9,600) and HET (slope = 28,000 ± 9,400) in particulate samples (Table S2). Quantification slopes for cobalamins after normalization with heavy internal standard showed little variation (Table S2, Fig S2).

**Figure 2:**
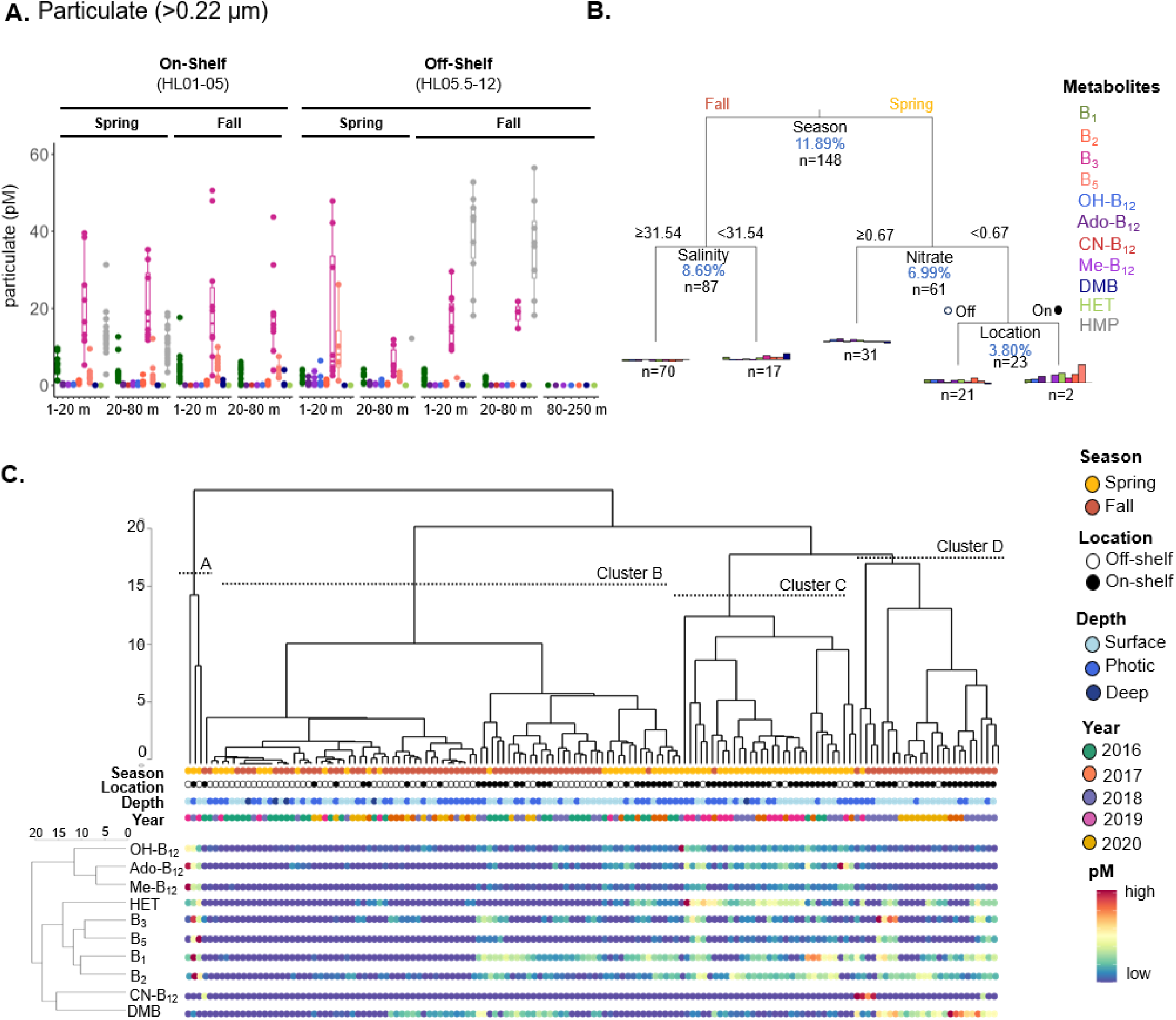
Measurements, explanatory variables, and trends in particulate b-vitamins and vitamers. (A) Particulate concentration (pM) of b-vitamins quantified in samples over spring and fall, on (HL01-HL05) and off (HL05.5-HL12) the Scotian Shelf in different depth bins (1 – 20 m, 20 – 80 m, 80 – 250 m). All years combined. (B) Multivariate regression tree (MRT) of particulate samples with a model R^2^ of 31.40%. The percent of variance explained by each factor, for the entire model, is indicated in blue along with the number of samples grouped at that split. Barplots represent metabolite concentrations scaled around the mean in the corresponding sample group. (C) Hierarchical clustering of samples based on particulate metabolite concentrations with associated season, location, depth, year, and relative particulate concentration of each metabolite respectively (high = red, low = blue). Four clusters (A, B, C, D) of samples are presented. Relative concentrations of metabolites are ordered based on hierarchical clustering (A, left dendrogram).

**Table 1:** Average particulate concentrations of metabolites, prevalence in samples, and ratios to carbon from the Halifax Line. Average limit of detection (LOD) in pM ± SD and R^2^ ± SD of calibration curves of four independent matrix groups are shown. Matrix group specific LOD and R^2^ is provided in Table S2. Prevalence represents the percentage of samples (%) the analyte was detected in (out of 148).

| analyte | limit of detection (pM) | R<sup>2</sup> | prevalence in samples (out of n=148) | average pM ± SD | average molar ratio to carbon ± SD (x 10<sup>-8</sup>) (1 m) |
| --- | --- | --- | --- | --- | --- |
| Me-B<sub>12</sub> | 0.05 ± 0.04 | 0.96 ± 0.03 | 53% | 0.17 ± 0.26 | 1.1 ± 0.79 |
| Ado-B<sub>12</sub> | 0.11 ± 0.08 | 0.94 ± 0.06 | 41% | 0.17 ± 0.18 | 1.1 ± 0.78 |
| OH-B<sub>12</sub> | 0.05 ± 0.02 | 0.98 ± 0.01 | 76% | 0.15 ± 0.19 | 1.8 ± 2.0 |
| CN-B<sub>12</sub> | 0.03 ± 0.02 | 0.97 ± 0.02 | 6% | 0.06 ± 0.03 | 0.87 ± 0.51 |
| DMB | 0.05 ± 0.02 | 0.98 ± 0.01 | 55% | 0.13 ± 0.11 | 4.0 ± 5.0 |
| B<sub>1</sub> | 2.33 ± 0.51 | 0.89 ± 0.08 | 57% | 3.4 ± 2.2 | 49 ± 47 |
| HET | 0.01 ± 0.01 | 0.99 ± 0.00 | 43% | 0.02 ± 0.01 | 0.25 ± 0.28 |
| B<sub>2</sub> | 0.24 ± 0.05 | 0.98 ± 0.02 | 71% | 0.53 ± 0.42 | 12 ± 15 |
| B<sub>5</sub> | 2.58 ± 0.96 | 0.96 ± 0.04 | 30% | 3.8 ± 4.3 | 2.8 ± 2.1 |
| B<sub>3</sub> | 7.95 ± 2.68 | 0.91 ± 0.08 | 45% | 25 ± 25 | 290 ± 240 |
| HMP* | 20.30 ± 14.70 | 0.82 ± 0.08 | 36% | 21 ± 14 | 810 ± 1300 |

\*Calibration curves for HMP were only reliable in spring on-shelf and fall off-shelf matrix groups

In particulate samples, OH-B_12_ and B_2_ were detected the most frequently while CN-B_12_ and B_5_ were only detected in 6% and 30% of samples respectively. No analyte was detected in 100% of the samples. Total B_12_ (Ado-, Me-, OH- and CN-B_12_ combined) ranged from 0.01 up to 4.7 pM over season, depth, and location. DMB, the lower ligand of B_12_, was detected in 55% of samples at a concentration of 0.13 ± 0.11 pM. B_1_ was detected in 57% of samples and ranged 3.4 ± 2.2 pM. HET ranged from 0.02 ± 0.01 pM and HMP was only reliably measured in two matrix groupings at a mean concentration of 21 ± 14 pM. Mean particulate B_2_ was 0.53 ± 0.42 pM and it was quantified in 71% of samples, while B_5_ was measured on average at 3.8 ± 4.3 pM. Particulate b-vitamin stoichiometry (mole b-vitamin to mole carbon), or molar ratios was calculated when possible (1 m depth, select stations) to account for differences in biomass across samples, and ranged from 10^-6^ to 10^-9^.

A multivariate regression tree (MRT) analysis (R^2^ = 31.40%) was performed to assess which factors best described the variance of the particulate metabolite samples (n = 148) (Fig 2B). For particulate metabolites, season explained the most variance (11.89%), shown as the first node of the tree. Alternative variables that could have been the first node are reported in Table S3. Samples from fall were split based on salinity, which explained 8.69% of the variance. Samples with ≥ 31.54 salinity (n = 70) had low concentrations of all measured metabolites. Samples with < 31.54 salinity (n = 17) in fall had higher concentrations of B_2_, B_3_, B_5_, and DMB. Samples collected in spring further split by nitrate (≥ and < 0.67 mmol m^-3^) which explained 6.99% of the total variance.

Samples with higher concentrations of nitrate (n = 31) had lower relative concentrations of B_2_, B_3_, and B_5_. Samples with lower concentrations of nitrate were then split by location (3.80% of total variance). Within this high nitrate in spring group, samples from on-shelf (n = 2) had higher concentrations of most metabolites. In general, particulate samples in spring had higher concentrations of OH-, Ado-, and Me-B_12_ and lower relative concentrations of DMB. Notably, while year and time of day were included in the MRT, they were not selected at any node.

Four clusters were chosen from the hierarchical clustering analysis based on the similarity of particulate metabolite concentrations in the samples (Fig 2C). The first cluster (A) contained three spring samples that had distinctly high concentrations of either Ado and Me-B_12_ or B_3_, B_5_, B_1_, or B_2_. Cluster B was the largest group containing samples from both seasons and locations that had relatively low concentrations of all metabolites with the exception of B_1_ and DMB. Cluster C was mainly comprised of on-shelf samples collected in spring which contained the highest observed concentrations of Ado and Me-B_12_, and HET. The majority of on-shelf samples from fall were found in Cluster D, which all had higher DMB, CN-B_12_, and B_3_. concentrations of all B_12_ forms. B-vitamin concentrations of samples are ordered based on hierarchical clustering (Fig 2C, left dendrogram) and generally grouped into 3 clusters. A grouping of Ado, Me and OH- B_12_, a grouping of HET, B_3_, B_5_, B_1_ and B_2_ and a grouping of CN-B_12_ and DMB.

#### Season-specific correlations in particulate samples

A season-specific Spearman’s rank correlation analysis of particulate metabolites and a suite of associated measurements, including pigments, in particulate samples from 1 m depth was performed (Fig 3A,B). Broadly, the strength and direction of correlation between variables and metabolites are different between seasons. Many metabolites, with the exception of B_5_ and CN-B_12_, correlated positively together in spring but these trends were not always mirrored in fall. Correlations between particulate organic carbon (POC) and chl *a* concentrations are season specific: no correlation is apparent in fall, but a strong positive correlation is seen in spring (Fig 3C). B_1_ and B_3_ correlated significantly (p-value < 0.05) with particulate organic carbon in fall but not spring (Fig 3D, E). As outlined by the green boxes, significant (p-value < 0.05) correlations of B_2_ and DMB with chl *a* were observed in fall that were absent in spring (Fig 3F, G).

**Figure 3:**
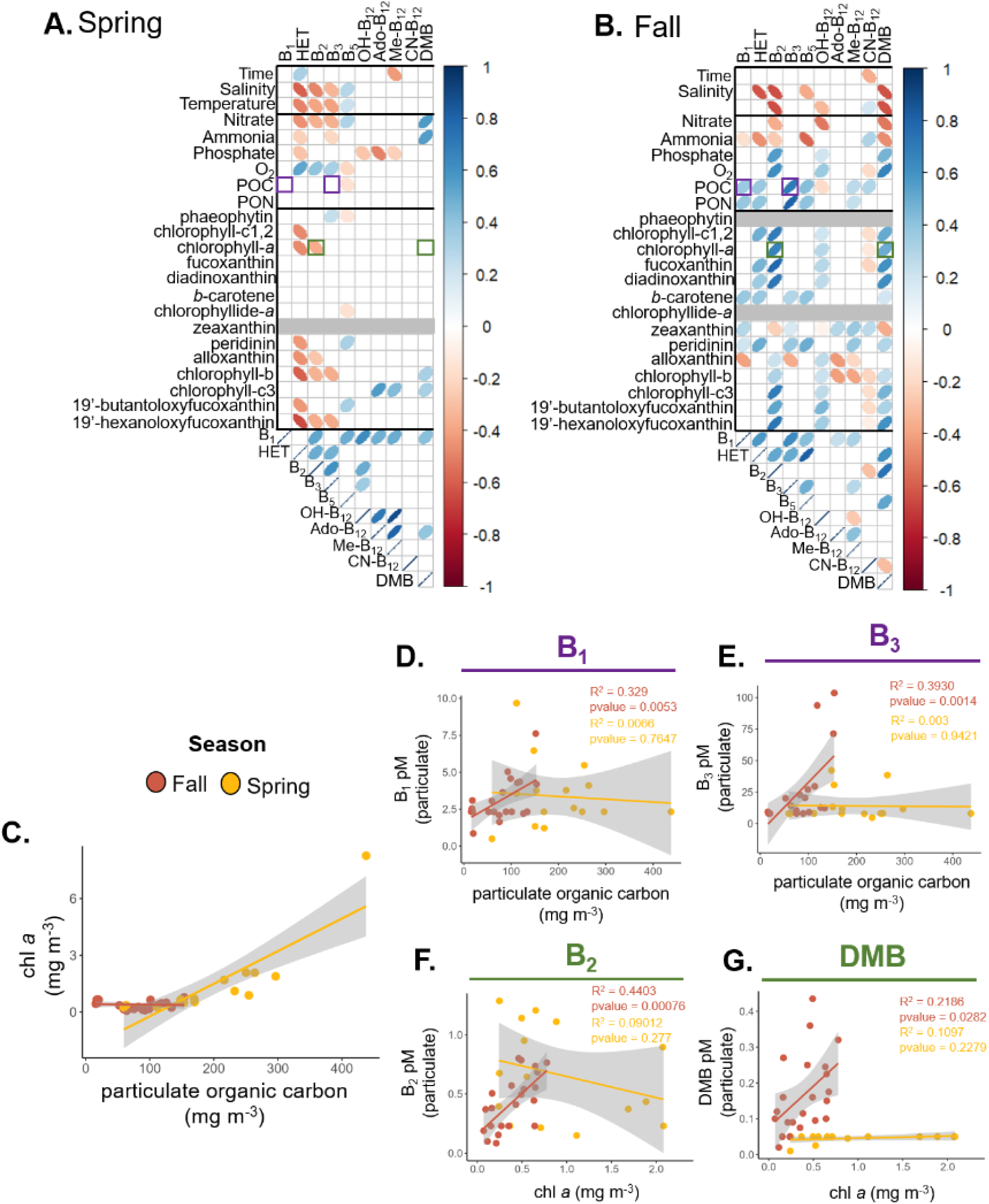
Season-specific correlations of environmental parameters with particulate b- vitamins and vitamers. Spearman rank correlations of environmental parameters (time, salinity, temperature, chl *a*), nutrient measurements (nitrate, ammonia, phosphate, O_2_, particulate organic carbon (POC), particulate organic nitrogen (PON)), pigment concentrations quantified by HPLC, and metabolites from particulate samples at 1 m depth in (A) spring and (B) fall. Bold lines separate groups of measurements. Grey boxes indicate that the HPLC pigment was not detected in that season. Green and purple boxes indicate relationships shown in (D-G). (C) Chl *a* concentrations (mg m^-3^) versus particulate organic carbon (mg m^-3^) in spring (yellow) and fall (orange). Particulate concentrations (pM) of (D) B_1_ and (E) B_3_ versus particulate organic carbon (mg m^-3^) in spring (yellow) and fall (orange). Particulate concentration (pM) of (F) B_2_ and (G) DMB versus chl *a* concentration (mg m^-3^) in spring (yellow) and fall (orange). R^2^ and p-values of the linear relationship of the plots are shown. Grey band represents the 95% confidence intervals of linear regression.

#### Dissolved samples

A total of 133 samples were extracted in duplicate for this study (Table 2, Fig 4). The average of the duplicated extractions from each sample is presented. Concentrations of 9 metabolites (OH-B_12_, CN-B_12_, DMB, HET, HMP, FAMP. B_2_, B_3_, B_6_) had reliable enough percent recovery to be quantified in dissolved samples (Fig 4A). Percent recovery of OH- and CN-B_12_ were >83% while recoveries for B_3_ and B_6_ were ∼53% and ∼47% respectively. Due to the selectivity of SPE, fewer metabolites were detected in dissolved phase (n = 9) than particulate (n =11) phase. Matrix effects, as determined from variations in calibration curve slopes between matrix groupings, were less evident in dissolved samples compared to particulate for analytes targeted in this study (Table S4).

**Figure 4:**
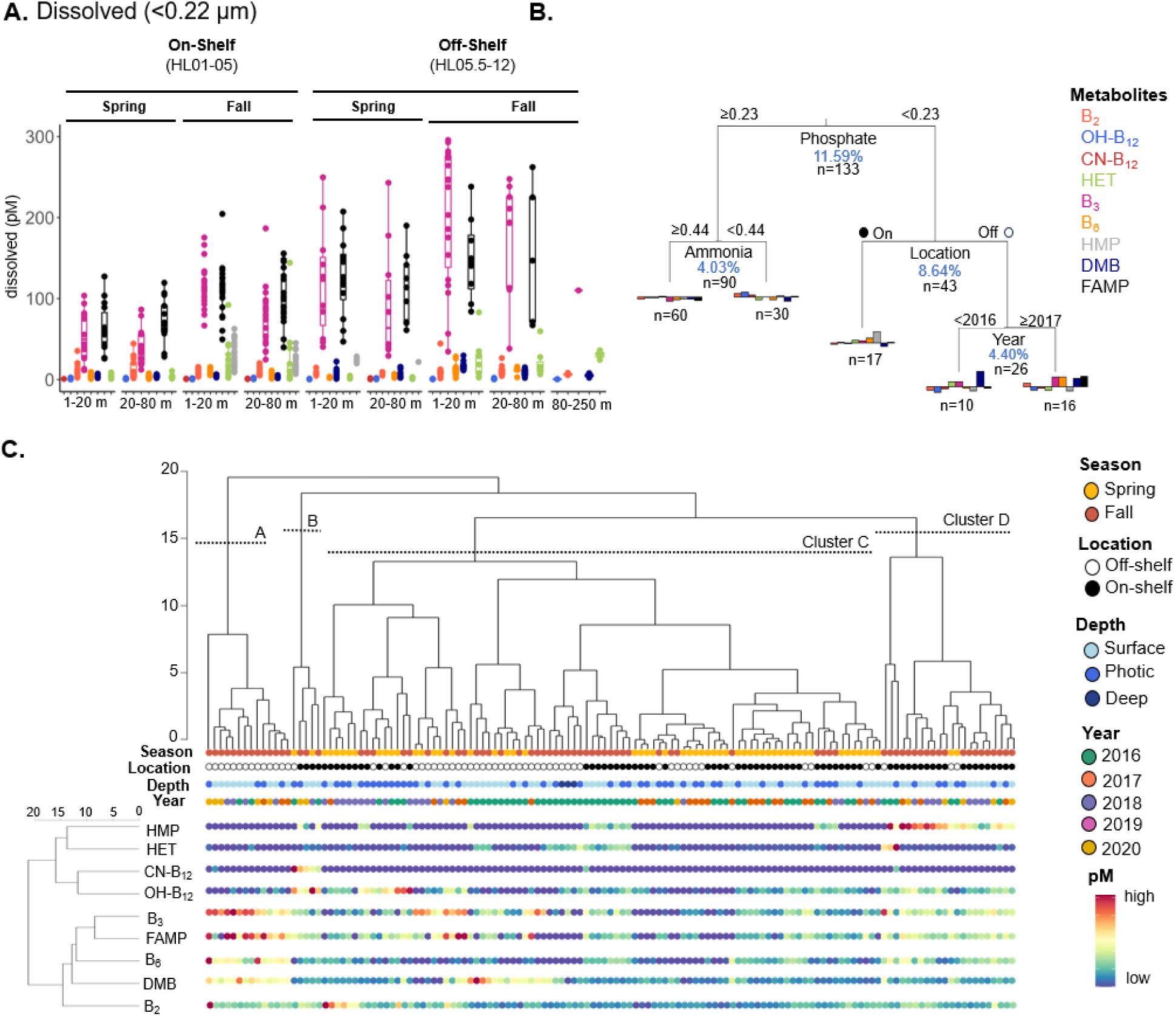
Measurements, explanatory variables and trends in dissolved b-vitamins and vitamers. (A) Dissolved concentration (pM) of b-vitamins quantified in samples over spring and fall, on (HL01-HL05) and off (HL05.5-HL12) the Scotian Shelf in different depth bins (1 – 20 m, 20 – 80 m, 80 – 250 m). All years combined. (B) Multivariate regression tree (MRT) of dissolved samples with an R^2^ of 28.7%. The percent of variability explained by each variable, for the entire model, is indicated in blue along with the number of samples grouped at that split. Barplots represent metabolite concentrations scaled to around the mean in the corresponding sample group. (C) Hierarchical clustering of samples based on dissolved metabolite concentrations with associated season, location, depth, year, and relative particulate concentration of each metabolite respectively (high = red, low = blue). Four clusters (A, B, C, D) of samples are presented. Relative concentrations of dissolved metabolites are ordered based on hierarchical clustering (A, left dendrogram).

**Table 2:** Average dissolved concentrations of analytes and prevalence in samples from the Halifax Line. Average percent recovery ± SD during solid phase extraction, limit of detection (LOD) in pM ± SD and R^2^ ± SD of calibration curves of four independent matrix groups. Matrix group-specific LOD and R^2^ is provided in Table S3. Prevalence represents the proportion of samples (%) the analyte was detected in (out of 148).

| compound | percent recovery (%) ± SD | limit of detection (pM) | R<sup>2</sup> | prevalence in of samples (of n=133) | average pM ± SD |
| --- | --- | --- | --- | --- | --- |
| OH-B<sub>12</sub> | 83 ± 19 | 0.35 ± 0.17 | 0.98 ± 0.01 | 75% | 0.61 ± 0.49 |
| CN-B<sub>12</sub> | 90 ± 18 | 0.73 ± 0.58 | 0.97 ± 0.03 | 6% | 0.78 ± 0.46 |
| DMB | 57 ± 25 | 4.6 ± 2.6 | 0.94 ± 0.05 | 95% | 6.9 ± 5.6 |
| HET | 75 ± 24 | 3.2 ± 3.4 | 0.95 ± 0.05 | 84% | 11 ± 20 |
| HMP | 84 ± 30 | 65 ± 11 | 0.95 ± 0.05 | 30% | 25 ± 13 |
| FAMP | 66 ± 17 | 150 ± 71 | 0.88 ± 0.07 | 79% | 130 ± 81 |
| B<sub>2</sub> | 75 ± 22 | 3.6 ± 1.7 | 0.97 ± 0.03 | 98% | 9.5 ± 7.0 |
| B<sub>3</sub> | 53 ± 27 | 140 ± 48 | 0.94 ± 0.05 | 86% | 110 ± 78 |
| B<sub>6</sub> | 47 ± 11 | 6.5 ± 2.4 | 0.93 ± 0.07 | 76% | 6.6 ± 4.9 |

OH-B_12_ was detected in 75% of dissolved samples at a mean concentration of 0.61 ± 0.49 pM. CN-B_12_ was measured in 6% of samples at 0.78 ± 0.46 pM. DMB was measured at 6.9 ± 5.6 pM in almost all dissolved samples (95%). HET and HMP were measured at 11 ± 20 and 25 ± 13 pM respectively. HMP was only detectable in 30% of samples while HET was detected in 84%. FAMP was measured at 79% of dissolved samples at 130 ± 81 pM. B_2_ was the most frequently detected dissolved metabolite (98% samples) and was found at 9.5 ± 7.0 pM. B_3_, in the form of niacin, was measured in 86% of samples at an average concentration of 110 ± 78 pM. B_6_ had an average of 6.6 ± 4.9 pM in 76% of samples.

A multivariate regression tree (MRT) analysis (R^2^ = 28.7%) was performed to assess which factors best described the variance of the dissolved metabolite samples (n=133) (Fig 4B). The variable that explained the most variance with the initial split of the entire dissolved metabolite dataset was phosphate (11.59%), with location being the next best alternative split for Node 1 (9.42%) (Table S5). Samples with higher concentrations of phosphate were then split by ammonia concentration which explained 4.03% of the model’s variance. Samples with ≥ 0.44 mmol m^-3^ ammonia had low concentrations of all dissolved metabolites except HET while samples < 0.44 mmol m^-3^ ammonia had higher concentrations of B_2_, OH-B_12_, and CN-B_12_. Samples with < 0.23 mmol m^-3^ phosphate were then split by location (on and off shelf) which explained 8.64% of the variance. Samples taken on-shelf had elevated concentrations of B_3_, B_6_, and HMP. Samples collected from off-shelf were then split into two groups based on sample year (4.40% of the model’s variance). Higher concentrations of B_2_, B_6_, and FAMP were measured in samples from 2017- 2020 compared to 2016.

Four distinct clusters were identified in the hierarchical clustering of dissolved metabolites (Fig 4C). The first group, cluster A, contained samples from fall, off-shelf with the highest concentrations of B_3_, FAMP, B_6_, and DMB. Cluster B was a group of on-shelf samples from both seasons where CN-B_12_ was measured. Cluster C was the largest group from both locations and seasons. Samples from fall, on-shelf group in cluster D and displayed the highest concentrations of HMP. Dissolved metabolites are ordered by hierarchal clustering (Fig 4C, left dendrogram). In dissolved samples, metabolites clustered into two distinct groups, the first of which contained CN-B_12_, OH- B_12_, HMP, and HET and the second cluster contained B_3_, FAMP, B_6_, DMB and B_2_.

#### Phase partitioning and evidence of cobalamin co-limitation

Five metabolites were reliably measured in both phases (HET, DMB, B_2_, B_3_, B_12_) and therefore could be used to investigate the phase partitioning (Fig 5A). DMB, HET, B_3_, and B_2_ were consistently enriched in dissolved phase compared to particulate phase (Fig 5A). B_12_ (particulate = Ado-, Me-, CN-, and OH-B_12_ combined) was enriched in the particulate phase, indicating concentration in cells was similar or more than the concentration in seawater (dissolved = CN- and OH-B_12_ combined), in spring samples. The concentration of OH-B_12_ in the dissolved phase over time shown as days since peak bloom, as defined in Fig 1 via remote sensing analysis, at 1 m depth, shows a general decline over time in the spring, and no sign of decline in the fall.

**Figure 5:**
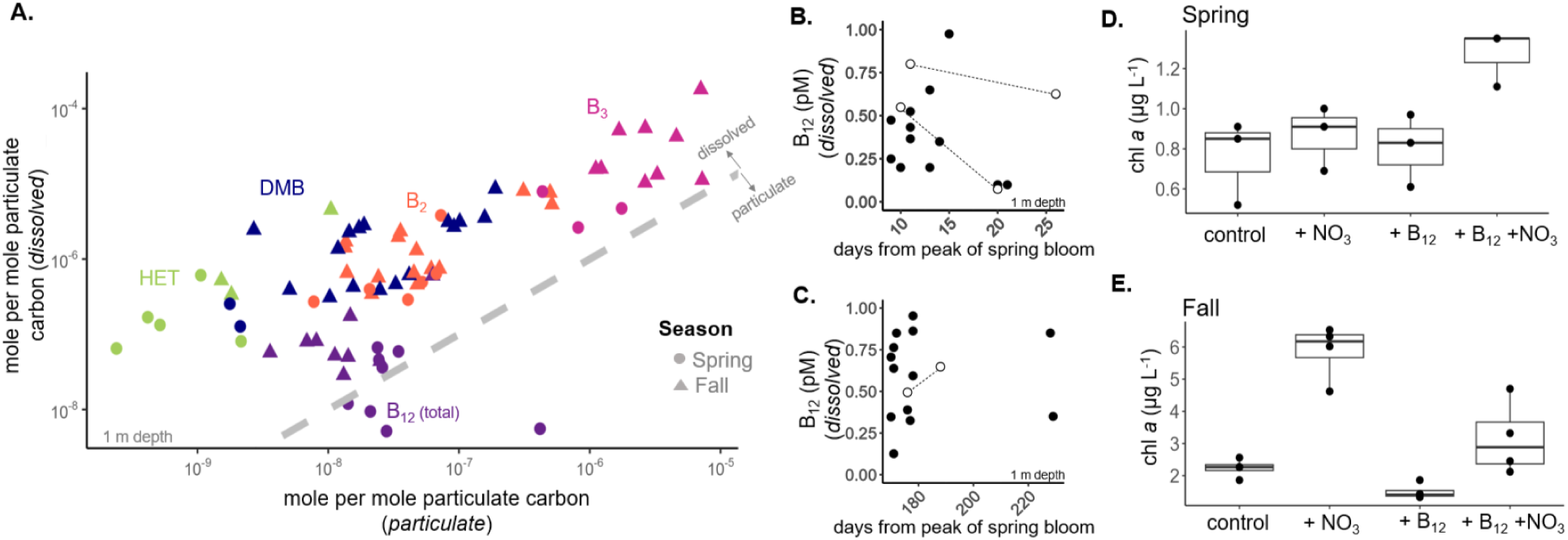
Phase partitioning of b-vitamins and vitamers, springtime B_12_ drawdown in dissolved phase and evidence of co-limitation. (A) Log_10_ scale of dissolved (Y-axis) versus particulate (X-axis) phase mole metabolite per mole particulate organic carbon from samples collected at 1 m depth in spring (circle) and fall (triangle). Grey dashed line indicates 1:1 relationship of dissolved to particulate. Points to the left of this line indicate dissolved enrichment while points to the right are indicative of particulate enrichment. Concentration of dissolved OH-B_12_, days from peak spring bloom (Fig 1F) in spring (B) and fall (C) from 1 m depth, open circles connected by dashed line are repeat occupation of station HL02 during the same cruise. Nutrient addition bottle incubation assay in (D) spring and (E) fall. Chl *a* (µg L^-1^) measured four (spring) and (fall) days after addition of +NO_3_ (10 µM), +B_12_ (100 pM) and +NO_3_+B_12_ (10 µM, 100 pM) collected on (B) May 3rd, 2017, and (C) Oct 5th, 2016, from HL02. Statistical tests for chl *a* measurements from bottle incubation assay provided in Table S6.

Nutrient addition bottle incubation assays were performed in both spring (2017) and fall (2016) with water collected from 5 m depth at station HL02 to examine how nitrate and cobalamin additions impacted microbial growth (Fig 5D, E, Fig S3). In spring, chl *a* concentration was highest four days after the addition of B_12_ and nitrogen together compared to both the control and individual nutrient addition treatments (Fig 5D).

Although no significant difference was seen between +NO_3_ + B_12_ treatment and +NO_3_ treatment with the conservative Tukey’s test (p-value = 0.083) (Table S6) there was a significant difference observed via paired t-test (p-value = 0.028). The total number of total photosynthetic eukaryotic cells per mL was significantly higher +NO_3_ + B_12_ treatment compared to all others (Fig S3, Tukey’s p-values range = 0.0083 to 0.0459) driven by the significant increase in small (<3 µm) photosynthetic eukaryotic cells (Fig S3). This evidence suggests that the community was indeed co-limited by nitrogen and cobalamin in spring. Notably, there was a significant decrease in *Synechococcus* in +NO_3_ + B_12_ treatment compared to others and no significant differences in bacterial cell counts between +NO_3_, +B_12_, and +NO_3_+B_12_ additions (Fig S3). In fall, the addition of nitrogen alone significantly increased chl *a* concentration more than any other treatment (Fig 5E, Table S6).

## Discussion

Our analyses of b-vitamins and vitamers in the NWA across different stations, depths, seasons, and years allowed for an in-depth investigation into the distribution and spatiotemporal variability of these metabolites in a dynamic oceanographic region. Arriving at detailed conclusions about the interaction between specific microbial community members and b-vitamins remain difficult because the driving factors controlling b-vitamin abundance in either phase have not been characterized, even in cultures. However, our study leverages these data to document and understand the variability in b-vitamins and vitamers in the ocean, identify patterns in their relationships with other oceanographic drivers, and investigate cobalamin dynamics in the NWA.

### Quantification and key forms of b-vitamins

Comparing b-vitamin concentrations from this study with those reported in other regions allows us to begin to constrain the distribution and variability of both particulate and dissolved b-vitamins in the ocean. A table outlining the extremely sparse measurements of vitamins and vitamers from the last decade and how they compare to the ones reported here is provided in the supplement (particulate: Table S7, dissolved: Table S8). The concentrations measured in this study fall into the range previously reported. However, to our knowledge, we report the first measurements of B_3_ (form nicotinamide) and DMB from particulate samples collected *in situ*.

One critical step forward in marine vitamin ecology is identifying important and dominant forms of b-vitamins in both phases, a topic that has recently been highlighted for cobalamin (Bannon et al. 2024). For example, the active forms of B_1_ are phosphorylated version, TMP (thiamine monophosphate) and TDP (thiamine diphosphate) but measuring these metabolites using reverse phase chromatography remain difficult, and the majority of measurements of B_1_ in the ocean, including in this study, are of the unphosphorylated cofactor which makes drawing conclusions about B_1_ quotas and B_1_ related processes in cells difficult.

In addition, B_3_ plays in a critical role in cells but it’s distribution in the ocean and role in microbial communities remain unclear (Table S7, S8). Here, we measure two forms of B_3_, niacin and niacinamide. Niacin is the hydrolyzed form of niacinamide and is thus expected to be a dominant form of B_3_ in the water column; however, additional research is required to verify this assumption. B_3_, in the form niacin, has been measured in dissolved samples from the North Sea at a range spanning from 26 to 46 pM (Bruns et al. 2022) and ranged from 11 to 319 pM in this study. In particulate samples, we report the first measurements of B_3_ (form niacinamide) which ranged from ∼3 to 124 pM (Table 1). B_3_, in the form of *niacin*, has been estimated at 11.3 and 18.6 pM in particulate samples from the North Pacific Subtropical Gyre (Boysen et al. 2021). Although we tried to quantify niacin in particulate samples, the calibration curve was unsuccessful, suggesting that we are underestimating the concentration of bioavailable B_3_ in the particulate phase.

Finally, B_6_ is hypothesized to be required for up to 1.5% of all enzymatic reactions in prokaryotes (Percudani and Peracchi 2003) and can be deficient in some fish species (Huang et al. 2005). However, B_6_ comes in six different forms that are all bioactive and the dominant forms of this co-factor in either phase remains unknown. Particulate measurements of pyridoxal and pyridoxal phosphate have been reported elsewhere (Boysen et al. 2021) and measurements of pyriodoxine in dissolved phase are reported here. The inconsistent measurements of B_6_ forms suggest that all current measurements underestimate the total biologically relevant B_6_ concentration in the ocean, and that the variability or influence of B_6_ may be overlooked.

Measuring the full suite of b-vitamin related compounds will be challenging, particularly due to the need to optimize analytical methods in order to capture the structurally diverse forms of each b-vitamin. We expect, however, that the continued interest in understanding the roles of b-vitamins in microbial communities, combined with advancing analytical capabilities to study them, will drive future research into the diversity and dominant forms of b-vitamins, and abiotic factors that might influence them.

### Spatiotemporal trends in b-vitamins

#### Distinct seasonal patterns in particulate samples

Our results suggest that season was an important driver of particulate b-vitamin dynamics (Fig 2B, Table S3). Notably, the differences between seasons goes beyond a combined increase or decrease of all metabolites in one season or the other, and instead we see distinct seasonal patterns in metabolite concentrations (Fig 2B; barplots at leaves of MRT). Previous work in the region has clearly shown distinct seasonal microbial assemblages and physicochemical characteristics in the NWA, both of which help contextualize the difference in b-vitamin profiles reported here (Li et al. 2006; Zorz et al. 2019; Robicheau et al. 2022).

In spring, chl *a* correlates with particulate organic carbon which suggests that surface biomass is dominated by autotrophs. In contrast, fall has major contributions from heterotrophic, non-chlorophyll *a* containing groups (Fig 3C) along with contributions from smaller phytoplankton and cyanobacteria (Li et al. 2006; Robicheau et al. 2022). Here we identified significant correlations of both B_2_ and DMB with chl *a* in fall, suggesting that fall phytoplankton may be the primary drivers of B_2_ and DMB concentrations in these samples, or vice versa (Fig 3F, G). Additionally, concentrations of particulate B_1_ and B_3_ correlated positively with particulate organic carbon in the fall (Fig 3D, E), a proxy for total community biomass, potentially suggesting that these may be key currencies in ‘recycler’ communities with relatively low new production compared to spring. Alternatively, this could suggest that b-vitamins quotas are higher in low-nitrogen environments. It remains unclear why correlations were absent in spring, though we do expect that variability in b-vitamin quotas over the course of the spring bloom could be quite high, given the large changes in phytoplankton physiology during the different stages of a bloom. Given that our sampling occurred at a range of stages of the spring bloom, changes in quotas could have prevented us from identifying any correlations with specific functional groups of microbes. Additional research into species-specific vitamin and vitamer quotas and factors that influence them are key to shedding light on the trends observed here.

#### No seasonally distinct patterns in dissolved concentrations

While metabolites in the dissolved phase are impacted by biological processes (e.g. production, excretion, lysis), this pool differs from the particulate phase because of the strong, direct impact of abiotic processes in the water column (e.g., hydrolysis and photolysis). Contrary to particulate samples, the two splits of dissolved b-vitamins that explained the most variance were nutrients (phosphate) and location (Fig 4B, Table S5).

In off-shelf samples we saw an increase of FAMP (Fig 4C), a degradation product of B_1_, which could indicate increased B_1_ degradation and/or limited FAMP uptake by the microbial community (Paerl et al, 2023). We see a similar trend with both DMB and B_3_ (niacin), which could be linked to enhanced light in the off-shelf samples due to lower amounts of dissolved organic matter blocking light penetration compared to on-shelf stations, however the driving factor of this accumulation is still unclear. The presence of other b-vitamins or reactive-oxygen species may also increase degradation or transformation of b-vitamins (Manzanares and Hardy 2010), and the rate of degradation relative to biological uptake or production for most of the b-vitamins remains unknown.

B-vitamins in low concentration in dissolved phase likely suggests quick degradation and/or biological uptake coupled with low production. Further work to describe the rates of b-vitamin degradation, uptake, and production is essential to provide context for interpreting dissolved b-vitamin measurements in the ocean and better predicting which of these vitamins is likely to exert controlling influences over microbial communities.

#### No discernable influence of time of day

Significant attention has been given to the influence of time of day and diel cycling on standing stocks of marine metabolites. A study from the North Pacific Subtropical Gyre highlighted that 70% of the measured particulate metabolites exhibited 24-hour periodicity, including B_5_, B_2_, and B_3_ (form niacin) and some osmolytes (Boysen et al. 2021). Furthermore, concentrations of both B_1_ and B_6_ (Sapelo Island, Georgia, USA) were higher in the seawater samples collected during the day (Gómez-Consarnau et al. 2018) and dissolved B_2_ was shown to be influenced by light in samples from the Sargasso Sea (Johnson et al. 2023). Such studies emphasize the light-regulated control on the microbial community and rapid response to their environment. Here, a relationship between B_2_ and time of day in dissolved or particulate phase was not observed across our datasets despite even sampling throughout the day and night (Fig S6). Furthermore, although time of day wass included in the MRT analysis, it did not come out as an main variable to split the data at any node (Fig 2B, 4B). Our results suggest that the variation contributed by time of day was overshadowed by other variables. Nevertheless, investigating the cycling of these metabolites at a higher resolution at one station would provide further insight into any daily patterns in vitamin production and loss in the NWA.

### B-vitamin variability within and between phases

#### Between different b-vitamins

B-vitamins support a wide range of processes in cells, with some required in high amounts and others required in low amounts. As such, we expect and observe that the range of requirements and concentrations in the ocean are wide as well. For example, B_3_ is consistently at higher concentrations than B_12_ in both phases in our data (Fig 2A, Table 1). B_3_ is the essential precursor of NAD/NADH or NADP/NADPH couples which are central to cellular processes (Kirkland and Meyer-Ficca, 2015) while B_12_ supports a small subset of specific enzymes, use patterns that are consistent with their high and low particulate concentrations, respectively. Half-saturation constants (Ks) are a useful metric to contextualize the range of b-vitamin concentrations at which auxotrophs may experience stress due to vitamin deprivation. For example, Ks values for B_12_ range from 0.02 to 13.1 pM for B_12_ in various phytoplankton (Tang et al, 2010) and have been document to be 37 nM and 4.6 µM for B_3_ in two *Rhodobacterales* isolates (Gregor et al, 2024). The reported B_3_ Ks values are orders of magnitude higher than the picomolar concentrations measured here (Table 2) which suggests that although B_3_ is present in relatively high concentrations, its availability may be impacting auxotrophs. B_3_ auxotrophy has been reported in B_12_-producing *Rhodobacteraceae* (Cooper et al. 2019) and particle associated bacterial isolates (Gregor et al, 2024), but a systematic investigation into B_3_ auxotrophy in marine microbial communities or an examination of the influence of B_3_ availability on B_12_ production have not yet been performed. Future studies to obtain additional Ks values for key molecules are essential to link analytical measurements and microbial ecology. Additional research into the distribution and variability of b-vitamin requirements, and factors that influence them, is essential for understanding the impact of these molecules on communities and processes.

#### Between b-vitamin and their vitamers

We observe large differences in concentrations of b-vitamins and their associated vitamers which may be consequential for their ecological roles. For example, various organisms that are auxotrophic for B_1_, like *E. huxleyi* (McRose et al, 2014), SAR11 (Carini et al, 2014) and other picoeukaryotic phytoplankton (Paerl et al, 2017) are able to grow on HMP, the pyrimidine part of the B_1_ molecule, alone. Here we show that particulate concentrations of HMP (2.88 – 56.50 pM) exceeded that of B_1_ (0.41 – 12.67 pM), both of which are far more abundant that HET, the thiazole part of the B_1_ molecule (0.01 – 0.06 pM) (Fig 2A, Table 1, Table S6). Although we were unable to reliably measure B_1_ in dissolved samples with the methods employed here, HMP and HET were measured consistently in dissolved samples at 6.9 – 54 pM and 0.44 – 144 pM respectively, both of which are within the range of B_1_ concentrations previously reported in literature (see Table S7). This work further uncovers the relative concentration of B_1_ and it’s vitamers in the ocean, highlighting the availability of HMP and its potential to be used to satisfy B_1_ demands.

### Seasonally distinct patterns in cobalamin influence

#### Cobalamin and nitrogen co-limitation in the spring

Cobalamin and nitrogen co-limitation has previously been described in various regions of the ocean, but this is the first evidence from the NWA and in relation to the spring bloom (Fig 5D, Fig S3). Comparisons between particulate and dissolved elemental stoichiometry of macronutrients and trace metals have been offered as an approach to potentially identify limiting nutrients, but has yet to be employed for vitamins (Moore et al, 2013). Indeed, we see that like nitrogen, cobalamin is enriched in particulate phase during an instance of bioassay-confirmed co-limitation (Fig 5A, D).

When we examined the concentration of dissolved cobalamin over time, in relation to the spring bloom, we observed a general decline in cobalamin availability at bloom termination, consistent with the idea that cobalamin co-limitation is playing a role in spring bloom demise (Fig 5B). A systematic investigation of cobalamin dynamics throughout the development and demise of algal blooms is a high priority for gaining further insight into control cobalamin may exert during these important events.

#### Evidence for cobalamin remodeling in the fall

Contrary to spring, cobalamin did not limit the microbial community in fall (Fig 5D). Concentrations of cobalamin in the dissolved phase were consistently elevated throughout the fall (Fig 5C). We hypothesize that this consistent cobalamin availability is due in part to an important role for cobalamin remodeling and recycling in the fall.

Several lines of evidence support this hypothesis. We observed increased concentrations of particulate DMB in fall compared to spring (Fig 2C). DMB is the lower ligand of cobalamin, required for remodeling cobalamin related compounds (cobinamides, pseudocobalamin) to generate cobalamin (Helliwell et al, 2016, Wienhausen et al, 2024). Although pseudocobalamin was not investigated here, it was previously observed up to 0.08 pM in the particulate phase in this region during fall (Bannon et al, 2024). Additionally, a previous metagenomics study in the region reported that cobalamin remodeling potential was elevated in fall compared to spring and this potential was mainly attributed to Alteromonadales, Pseudomonadales, Rhizobiales, Oceanospirilalles, Rhodobacteraceae, and Verrucomicrobia (Soto et al, 2023). We suggest that the increased DMB measured here, combined with the availability of psB_12_ and increased presence of organisms with remodeling activity in the fall, is evidence that cobalamin remodeling plays an important role in cobalamin dynamics on the NWA during this season, contributing to its consistent availability in the dissolved phase. Future work investigating remodeling rates, precursors, and key players would help clarify the overall impact remodeling has on cobalamin dynamics.

#### Seasonal vitamin dynamics in the NWA

A picture of seasonal differences in the roles of different vitamins in this NWA system is emerging, providing further insight into the role of different processes in vitamin supply and demand and their ultimate impact on ecosystems. For cobalamin, these data are consistent with cobalamin supply to the surface ocean at the beginning of a growing season being influenced by delivery, via mixing processes, of cobalamin produced at depth (Menzel and Spaeth 1962), which is then drawn down throughout the course of the spring bloom. Toward the end of the bloom, when cobalamin co-limitation is present, intimate associations between cobalamin producers and consumers may be key to continued cobalamin use by auxotrophs and facultative consumers. In contrast, cobalamin remodeling, recycling and regeneration in the oligotrophic and heterotroph- dominated fall appears to play a key role in maintaining elevated dissolved cobalamin levels, when cobalamin demand in key community members may also be satisfied through bacterivory or predation. This emerging picture has many parallels in macronutrient cycling that should be explored. Continued focus on quantifying the role of each of these processes in cobalamin production, use, and transformations, has the potential to generate new frameworks that can be leveraged to conceptually and numerically model vitamin impacts on marine ecosystems.

## Supporting information

Manuscript Data

## Author contribution statement

**CB:** Conceptualization, Methodology, Investigation, Formal Analysis, Visualization, Data Curation, Writing – Original Draft, Writing – Review & Editing.

**PLW:** Methodology, Investigation, Formal Analysis, Writing – Review & Editing.

**ER:** Methodology, Investigation, Formal Analysis, Writing – Review & Editing.

**KJM:** Investigation, Writing – Review & Editing

**AG:** Investigation, Writing – Review & Editing

**MR:** Investigation, Writing – Review & Editing

**ED:** Resources, Writing – Review & Editing.

**LB:** Resources, Writing – Review & Editing.

**JL:** Funding Acquisition, Resources, Writing – Review & Editing.

**EMB:** Conceptualization, Methodology, Supervision, Funding Acquisition, Resources, Writing – Original Draft, Writing – Review & Editing.

## Acknowledgements

We are grateful to the Fisheries and Oceans Canada’s Atlantic Zone Monitoring Program (AZMP) based at the Bedford Institute of Oceanography for providing the opportunity for the ship-based collection of samples analyzed in this study, and to Zoe Finkel and Anitra Ingalls for helpful discussions. This study was funded by Ocean Frontier Institute (Module C and NWA-BCP), NSERC Discovery Grant RGPIN2015- 05009 to EMB, Simons Foundation Grant 504183 to EMB, Simons Foundation CBIOMES Award ID 1001702 to EMB, Canada Research Chair Support to EMB, and NSERC CGS-D to CB.

