## Supplement Info (Table S1-S8 and Fig S1-S4) for "Seasonal patterns in b-vitamins and cobalamin co-limitation in the Northwest Atlantic"

### Supporting Information

**Document Includes:**

***Supporting methods***

Analytical standards

**Tables S1-**

Table S1: Transition list for targeted analytes
Table S2: Matrix group specific limit of detection and quantification for particulate samples.
Table S3: MRT analysis output (node 1) indicating key explanatory variables of particulate metabolite data set.
Table S4: Matrix group specific limit of detection and quantification for dissolved samples.
Table S5: MRT analysis output (node 1) indicating key explanatory variables of Dissolved metabolite data set.
Table S6: Statistical tests for chlorophyll a results from bottle incubation assays.
Table S7: Literature review of key particulate environmental measures of b-vitamin and vitamer since 2014.
Table S8: Literature review of key dissolved environmental measures of b-vitamin and vitamer since 2014.

**Figures S1-6**

Figure S1: Matrix grouping experimental analysis set up for Acclaim C18 column.
Figure S2: Calibration curves of Me-, Ado-, CN-, and OH-$B_{12}$ in particulate samples run on Acclaim C18 column before and after normalizing with heavy-CN-$B_{12}$ internal standard.
Figure S3: Flow cytometry data for nutrient addition bottle incubation assay in Spring, 2017 at HL02.
Figure S4: Dissolved and particulate concentrations (pM) of $B_2$ over time.

***Supplemental Methods***
**Analytical standards**

Metabolite standards CN-$B_{12}$ (≥98%, Fisher BioReagents), Me-$B_{12}$ (≥97%, Sigma-Aldrich), Ado-$B_{12}$ (≥97%, Sigma-Aldrich), OH-$B_{12}$ (≥95%, Supelco), DMB (5,6-dimethylbenzimidazole) (≥99%, Sigma-Aldrich), $B_5$ (p-pantothenic acid) (≥98%, Sigma-Aldrich), SAM (80%, Sigma-Aldrich), $B_6$ (pyridoxine) (>90%, Sigma-Aldrich), $B_1$ (thiamin hydrochloride) (≥95%, Sigma-Aldrich), HET (4-methyl-5-thiazoleethanol) (≥95%, Sigma-Aldrich), HMP (4-amino-5-hydroxymethyl-2-methylpyrimidine) (>95%, Enamine Ltd), FAMP (>90%, Toronto Research Chemicals), $B_2$ (%, Sigma-Aldrich), $B_3$ (niacin) (>90%, Sigma-Aldrich) and $B_3$ (niacinamide) (≥99%, Sigma-Aldrich) were obtained and primary stock solutions were prepared by dissolving 1 mg of each compound in 1 mL of Optima LC–MS grade water and stored at -80 °C until use.

***Supplemental Tables***

**Table S1:** Selected reaction monitoring mass spectrometry parameters for metabolites measured in this study.

| Analyte | Name | **SRM (m/z) precursor** → product | Collision Energies (eV) | Retention time (min) |
|---|---|---|---|---|
| Me-$B_{12}$ | methyl-cobalamin | **673.5** → 972.5, 665.9, 359.1 | 25.2 | 4.1 |
| Ado-$B_{12}$ | ado-cobalamin | **790.9** → 972.5, 665.9, 359.1 | 29.2 | 3.9 |
| OH-$B_{12}$ | hydroxy-cobalamin | **665.0** → 913.4, 636.0, 359.1 | 24.9 | 3.6 |
| CN-$B_{12}$ | cyano-cobalamin | **678.4** → 997.5, 635.9, 359.1 | 25.4 | 3.7 |
| heavy-CN-$B_{12}$ | | **681.9** → 997.5, 639.4, 366.1 | 25.4 | 3.7 |
| DMB | 5,6-dimethylbenzimidazole | **147.2** → 132.1, 120.1, 104.1 | 25.9 | 3.8 |
| $B_1$ | thiamine | **265.2** → 144.2, 122.3, 113.3 | 15.8 | 2.5 |

| | | | | |
|---|---|---|---|---|
| HET | 4-methyl-5-thiazoleethanol | **144.1** → 113.2, 112.2, 80.3, 71.3 | 19.5 | 3.5 |
| HMP | 4-amino-5-hydroxymethyl-2-methylpyrimidine | **140.1** → 122.2, 54.3 | 18.7 | 2.2 |
| FAMP | N-formyl-4-amino-5-aminomethyl-2-methylpyrimidine | **167.1** → 122.1, 54.1 | 15 | 2.4 |
| $B_2$ | riboflavin | **377.2** → 243.1, 198.1, 172.1, 145.1 | 22 | 3.8 |
| $B_5$ | d-pantothenic acid | **220.1** → 116.0, 90.0 | 17 | 3.0 |
| $B_3$ | niacinamide | **123.1** → 96.1, 80.1, 78.1, 53.2 | 17 | 2.9 |
| $B_3$ | niacin | **124.1** → 106.1, 96.1, 80.1, 78.1, 53.2 | 17 | 2.5 |
| $B_6$ | pyridoxine | **170.1** → 134, 124.1, 106.1, 77.1 | 34 | 2.5 |

**Table S2**: Slope, limit of detection (fmol on analytical column and pM value) and calibration curve $R_2$ of metabolites quantified in this particulate samples in specific matrix groups based on season and station location. Cobalamins (identified with the *) have been normalized to internal standard of heavy cyanocobalamin as seen in Figure S2.

| Matrix Group | Analyte | Slope | LOD (fmol on analytical column) | LOD (pM) | $R^2$ |
|---|---|---|---|---|---|
| fall, off-shelf | *Me-$B_{12}$ | 0.23 | 0.15 | 0.009 | 0.92 |
| | *OH-$B_{12}$ | 0.36 | 0.47 | 0.028 | 0.96 |
| | *Ado-$B_{12}$ | 0.17 | 0.64 | 0.038 | 0.97 |
| | *CN-$B_{12}$ | 0.097 | 0.36 | 0.021 | 0.99 |
| | DMB | 41000 | 0.68 | 0.041 | 0.96 |
| | $B_1$ | 33 | 46 | 2.8 | 0.97 |
| | HET | 24000 | 0.17 | 0.01 | 0.98 |
| | HMP | 360 | 580 | 35 | 0.74 |
| | $B_2$ | 1900 | 2.7 | 0.16 | 0.94 |
| | $B_3$ (niacinamide) | 780 | 140 | 8.6 | 0.78 |
| | $B_5$ | 86 | 53 | 3.2 | 0.98 |
| fall, on-shelf | *Me-$B_{12}$ | 0.3 | 0.95 | 0.057 | 0.98 |
| | *OH-$B_{12}$ | 0.54 | 1.2 | 0.072 | 0.98 |
| | *Ado-$B_{12}$ | 0.29 | 4.2 | 0.25 | 0.84 |
| | *CN-$B_{12}$ | 0.15 | 0.95 | 0.057 | 0.93 |
| | DMB | 25000 | 1.4 | 0.082 | 0.99 |
| | $B_1$ | 21 | 47 | 2.8 | 0.77 |
| | HET | 15000 | 0.45 | 0.027 | 0.99 |
| | $B_2$ | 1600 | 3.7 | 0.22 | 0.99 |
| | $B_3$ (niacinamide) | 780 | 160 | 9.8 | 0.97 |

| | | | | | |
|---|---|---|---|---|---|
| | $B_5$ | 92 | 61 | 3.7 | 0.89 |
| spring, off-shelf | *Me-$B_{12}$ | 0.26 | 1.9 | 0.11 | 0.96 |
| | *OH-$B_{12}$ | 0.36 | 0.67 | 0.04 | 0.99 |
| | *Ado-$B_{12}$ | 0.28 | 1.6 | 0.097 | 0.99 |
| | *CN-$B_{12}$ | 0.11 | 0.53 | 0.031 | 0.99 |
| | DMB | 52000 | 0.63 | 0.038 | 0.99 |
| | $B_1$ | 160 | 27 | 1.6 | 0.89 |
| | HET | 41000 | 0.12 | 0.007 | 0.99 |
| | $B_2$ | 3100 | 5 | 0.3 | 0.99 |
| | $B_3$ (niacinamide) | 3000 | 57 | 3.4 | 0.97 |
| | $B_5$ | 200 | 37 | 2.2 | 0.96 |
| spring, on-shelf | *Me-$B_{12}$ | 0.29 | 0.58 | 0.035 | 0.99 |
| | *OH-$B_{12}$ | 0.35 | 0.95 | 0.057 | 0.99 |
| | *Ado-$B_{12}$ | 0.29 | 1.1 | 0.068 | 0.96 |
| | *CN-$B_{12}$ | 0.12 | 0.24 | 0.015 | 0.97 |
| | DMB | 41000 | 0.55 | 0.033 | 0.99 |
| | $B_1$ | 170 | 34 | 2.1 | 0.94 |
| | HET | 30000 | 0.17 | 0.01 | 0.99 |
| | HMP | 1500 | 93 | 5.6 | 0.9 |
| | $B_2$ | 4600 | 4.3 | 0.26 | 0.99 |
| | $B_3$ (niacinamide) | 3900 | 170 | 10 | 0.91 |
| | $B_5$ | 330 | 20 | 1.2 | 0.99 |

**Table S3:** Output for primary split (Node 1) of the multivariate regression tree analysis for particulate samples showing other variables that explain variability in entire data set and their optimal breakpoints.

| Variable | Threshold | Direction | Variability explained of entire data set |
|---|---|---|---|
| Season | *NA* | splits as left/right | 11.89% |
| Salinity | <32.30 | to the right | 10.40% |
| Temperature | <3.8284 | to the right | 9.17% |
| Location | *NA* | splits as left/right | 8.62% |
| Time | <2324 | to the left | 5.31% |

**Table S4:** Slope, limit of detection (fmol on analytical column and pM value) and calibration curve $R_2$ of analytes quantified in this dissolved samples in specific matrix groups based on season and station location.

| Matrix Group | Analyte | Slope | LOD (fmol on column) | LOD (pM) | $R^2$ |
|---|---|---|---|---|---|
| fall, off-shelf | OH-$B_{12}$ | 16000 | 0.48 | 0.6 | 0.99 |
| | CN-$B_{12}$ | 7700 | 0.8 | 0.9 | 0.99 |
| | DMB | 14000 | 4.7 | 8.2 | 0.95 |
| | HET | 20000 | 6.7 | 9 | 0.99 |
| | HMP | 230 | 61 | 72 | 0.94 |
| | FAMP | 1600 | 180 | 270 | 0.81 |
| | $B_2$ | 9500 | 4.9 | 6.5 | 0.99 |
| | $B_3$ (niacin) | 2400 | 110 | 200 | 0.87 |
| | $B_6$ | 31000 | 4.7 | 10 | 0.91 |
| fall, on-shelf | OH-$B_{12}$ | 18000 | 0.2 | 0.2 | 0.99 |
| | CN-$B_{12}$ | 8300 | 0.3 | 0.3 | 0.99 |
| | DMB | 20000 | 1.2 | 2.1 | 0.99 |
| | HET | 19000 | 0.9 | 1.2 | 0.99 |
| | HMP | 200 | 59 | 69 | 0.96 |
| | FAMP | 2100 | 68 | 100 | 0.93 |
| | $B_2$ | 10000 | 1.9 | 2.5 | 0.99 |
| | $B_3$ (niacin) | 1400 | 73 | 140 | 0.98 |
| | $B_6$ | 23000 | 2.9 | 6.1 | 0.99 |
| spring, off-shelf | OH-$B_{12}$ | 16000 | 0.33 | 0.4 | 0.97 |
| | CN-$B_{12}$ | 8400 | 1 | 1.6 | 0.92 |
| | DMB | 17000 | 3 | 6.1 | 0.95 |
| | HET | 22000 | 1 | 1.4 | 0.97 |
| | HMP | 260 | 61 | 72 | 0.99 |
| | FAMP | 1700 | 78 | 120 | 0.98 |
| | $B_2$ | 8000 | 2 | 2.6 | 0.96 |
| | $B_3$ (niacin) | 3200 | 75 | 140 | 0.99 |

| | | | | | |
|---|---|---|---|---|---|
| | $B_6$ | 30000 | 3 | 6.6 | 0.99 |
| spring, on-shelf | OH-$B_{12}$ | 15000 | 0.17 | 0.2 | 0.98 |
| | CN-$B_{12}$ | 8200 | 0.06 | 0.1 | 0.99 |
| | DMB | 24000 | 1 | 2.1 | 0.85 |
| | HET | 32000 | 1 | 1 | 0.86 |
| | HMP | 390 | 38 | 45 | 0.86 |
| | FAMP | 2300 | 66 | 100 | 0.81 |
| | $B_2$ | 8500 | 2 | 2.6 | 0.92 |
| | $B_3$ (niacin) | 3100 | 34 | 66 | 0.9 |
| | $B_6$ | 38000 | 1.5 | 3.2 | 0.82 |

**Table S5:** Output for primary split (Node 1) of the multivariate regression tree analysis for dissolved samples showing other variables that explain variability in entire data set and their optimal breakpoints.

| **Variable** | **Threshold** | **Direction** | **Variability explained of entire data set** |
|---|---|---|---|
| Phosphate | <0.232 | to the right | 11.59% |
| Location | *NA* | splits as left/right | 9.42% |
| Temperature | <19.029 °C | to the left | 8.90% |
| Nitrate | <0.3725 | to the right | 8.11% |
| Season | *NA* | splits as left/right | 7.88% |

**Table S6:** P-values from both Tukey's and unpaired t-tests for pairwise comparison of chlorophyll *a* concentration from bottle incubation assay

| Season | Pairwise Comparison | Adjusted p-value (Tukey test) | P-value (unpaired t-test) |
|---|---|---|---|
| Spring | Control vs $B_{12}$ | 0.991 | 0.813 |
| | $NO_3$ vs $B_{12}$ | 0.964 | 0.659 |
| | $NO_3$ + $B_{12}$ vs $B_{12}$ | 0.042 | 0.024 |
| | $NO_3$ vs Control | 0.872 | 0.517 |
| | $NO_3$ + $B_{12}$ vs Control | 0.029 | 0.024 |
| | $NO_3$ + $B_{12}$ vs $NO_3$ | 0.083 | 0.028 |
| Fall | Control vs $B_{12}$ | 0.521 | 0.008 |
| | $NO_3$ vs $B_{12}$ | 1.31E-05 | 0.0001 |
| | $NO_3$ + $B_{12}$ vs $B_{12}$ | 0.037 | 0.031 |
| | $NO_3$ vs Control | 8.27E-05 | 0.0002 |
| | $NO_3$ + $B_{12}$ vs Control | 0.351 | 0.176 |
| | $NO_3$ + $B_{12}$ vs $NO_3$ | 0.001 | 0.009 |

**Table S7:** Literature review of key particulate environmental measures of b-vitamin and vitamer since 2014. *Estimated pM concentration

| Paper | Location | $B_1$ (pM) | $B_2$ (pM) | Niacin ($B_3$) (pM) | Nicotinamide ($B_3$) (pM) | $B_5$ (pM) | Pyridoxine ($B_6$) (pM) | Pyridoxal ($B_6$) (pM) | DMB (pM) | HET (pM) | HMP (pM) | FAMP (pM) | OH- $B_{12}$ (pM) | Ado- $B_{12}$ (pM) | Me- $B_{12}$ (pM) | CN- $B_{12}$ (pM) | Me-ps$B_{12}$ (pM) | SAM (pM) |
|---|---|---|---|---|---|---|---|---|---|---|---|---|---|---|---|---|---|---|
| Bittner et al, 2024 | Roskilde fjord | 1.16 – 7.78 | | | | | | | | 0.01 – 0.54 | 2.21 – 41.42 | 0.35 – 2.86 | | | | | | |
| Johnson et al, 2023 | Atlantic Ocean | | 0.003 – 0.7 | | | | 0.1 – 2 | | | | | | | | | | | |
| Boysen et al, 2023* | North Pacific | | 0.217 | 44.1 | | 3.31 | | 4.68 | | | | | 0.095 | | | | | 43.9 |
| Suffridge et al, 2018 | Med Sea | 3.77 – 204.59 | | | | | | | | | 1.87 – 84.81 | | 1.21 – 19.20 | 0.07 – 6.86 | 0.03 – 6.89 | 0.01– 1.22 | | |
| Suffridge et al, 2017 | SPOT | 481 ± 204 | 129 ± 45.7 | | | | | | | | 40.8 ± 14.8 | | 40.8 ± 14.8 | 5.82 ± 6.53 | 3.14 ± 5.02 | 1.54 ± 2.46 | | |
| | Atlantic Station | 3.53 – 44.4 | | | | | | | | | 5.91– 45.56 | | 1.21 – 9.11 | 0.52 – 4.53 | 0.19 – 2.46 | <LOD – 1.21 | | |
| Heal et al, 2017 | North Pacific Ocean | | | | | | | | | | | | 0.01 – 0.24 | 0.00 – 0.05 | 0.00 – 0.02 | | 0.00- 0.09 | |
| Bannon et al, 2024 | Northwest Atlantic Ocean | | | | | | | | | | | | | | | | 0.01 – 0.08 | |
| This study | Northwest Atlantic Ocean | 0.41 – 13.00 | 0.09 –2.90 | NA | 2.50 – 120.00 | 0.60 – 26.00 | NA | NA | 0.01 – 0.52 | 0.01 – 0.06 | 2.90 – 56.00 | NA | 0.01 – 1.60 | 0.02 – 0.94 | 0.01 – 2.00 | 0.02 – 0.10 | NA | NA |

**Table S8:** Literature review of key dissolved environmental measurements of b-vitamin and vitamer since 2014. *Estimated based on plot.

| Paper | Location | Thiamine $B_1$ (pM)* | $B_2$ (pM) | Niacin ($B_3$) (pM) | Pyridoxine ($B_6$) (pM) | Pyridoxal ($B_6$) (pM) | OH-$B_{12}$ (pM) | CN-$B_{12}$ (pM) | DMB (pM) | HET (pM) | HMP (pM) | FAMP (pM) |
|---|---|---|---|---|---|---|---|---|---|---|---|---|
| Bittner et al, 2024 | Roskilde fjord | 37.97 – 195.83 | | | | | | | | 2.01 – 12.11 | 9.94 – 36.95 | 11.85 – 94.57 |
| Longnecker et al, 2024 | Northwester Sargasso Sea | 0 – 200* | 0 – 0.6* | | 0 – 1* | | | 0 – 0.2* | | | | |
| Johnson et al, 2023 | Atlantic Ocean | | 0.7 – 12 | | | | | | | | | |
| Paerl et al, 2023 | Scotian Shelf and Slope, NS, Canada | | | | | | | | | 6 –13 | 22 – 36 | 14 – 36 |
| | Neuse River Estuary, NC, USA | 43 – 83 | | | | | | | | 1 – 5 | 21 – 52 | 23 – 36 |
| Bruns et al, 2023 | North Sea | 5-50* | | ~20 | <10 | | | 10* | <5* | <5* | ~10* | |
| Bruns et al, 2022 | North Sea (SRM values) | 26.8 ± 7.5 | 35.9 ± 2.4 | 26.0 ± 2.4 (PRM) 46.3 ± 9.6 (full scan) | 31.3 – 91 | | | 15.4 ± 1.7 | 13.7 ± 0.8 | 1.4 ± 0.2 | 15.8 ± 2.4 | |
| Suffridge et al, 2020 | North Atlantic | 8.36 –353 | | | | | | | | 0.03 – 30.8 | 0.11 – 3.45 | |
| Gómez-Consarnau et al. 2018 | Sapelo Island, Georgia, USA | 0.5 – 5 | | | 0.1 – 8.9 | | | | | | | |
| Suffridge et al, 2018 | Med Sea | 0.49 – 253 | | | | | 0.09 – 3.13 | 0.02 – 0.08 | | | <LOD – 190 | |
| Suffridge et al, 2017 | Atlantic Station | 33.4– 457.1 | | | | | 3.1 ± 0.2 | 0.8 ± 0.4 | | | 2.5– 28.3 | |
| Cohen et al, 2017 | Pacific Ocean | 3.3 – 9.5 | 1.3 – 9 | | | 0.48 – 5.49 | | 0.18 – 0.25 | | | | |
| Carini et al, 2014 | Sargasso Sea | up to 23 | | | | | | | | | 2.5 – 35 | |
| Heal et al, 2014 | Hood Canal | 0.58 – 1.5 | 45 – 128 | | 1.3 – 5.7 | | 1.56 – 5.8 | <0.7 | | | | |
| This study | Northwest Atlantic Ocean | NA | 1.50 – 44.00 | 11.00 – 320.00 | 0.60 – 28.00 | NA | 0.10 – 2.50 | 0.02 – 1.70 | 0.80 – 30.00 | 0.40 – 140.00 | 6.90 – 54.00 | 26.00 – 350.00 |

***Supplemental Figures***

**Figure S1:** Experimental design for matrix-specific calibration curves and analysis. Particulate and dissolved samples were grouped based season and location. Calibration curves were made from pooled quality control (QC) sample. Matrix-group specific slope and limit of detection (LOD) were then applied to samples in that matrix grouping.

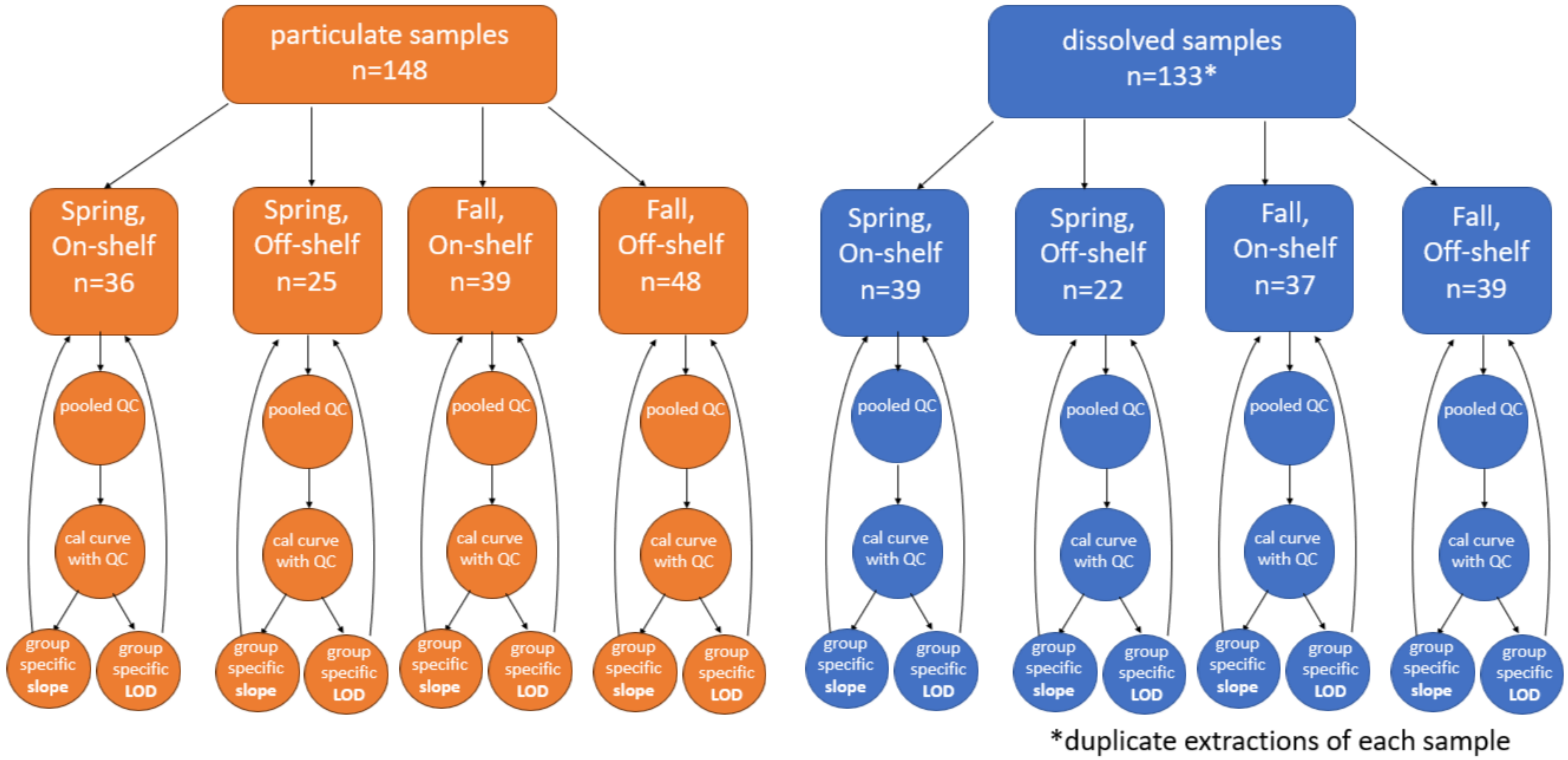

**Figure S2:** Matrix-group specific calibration curves reported in (A) peak area and (B) peak area normalized to heavy-CN-$B_{12}$ internal standard (IS) versus fmol on analytical column for all cobalamin forms (Ado, CN, Me, and OH-$B_{12}$) in particulate samples. Grey shading indicates the standard error confidence internal of linear model relationship.

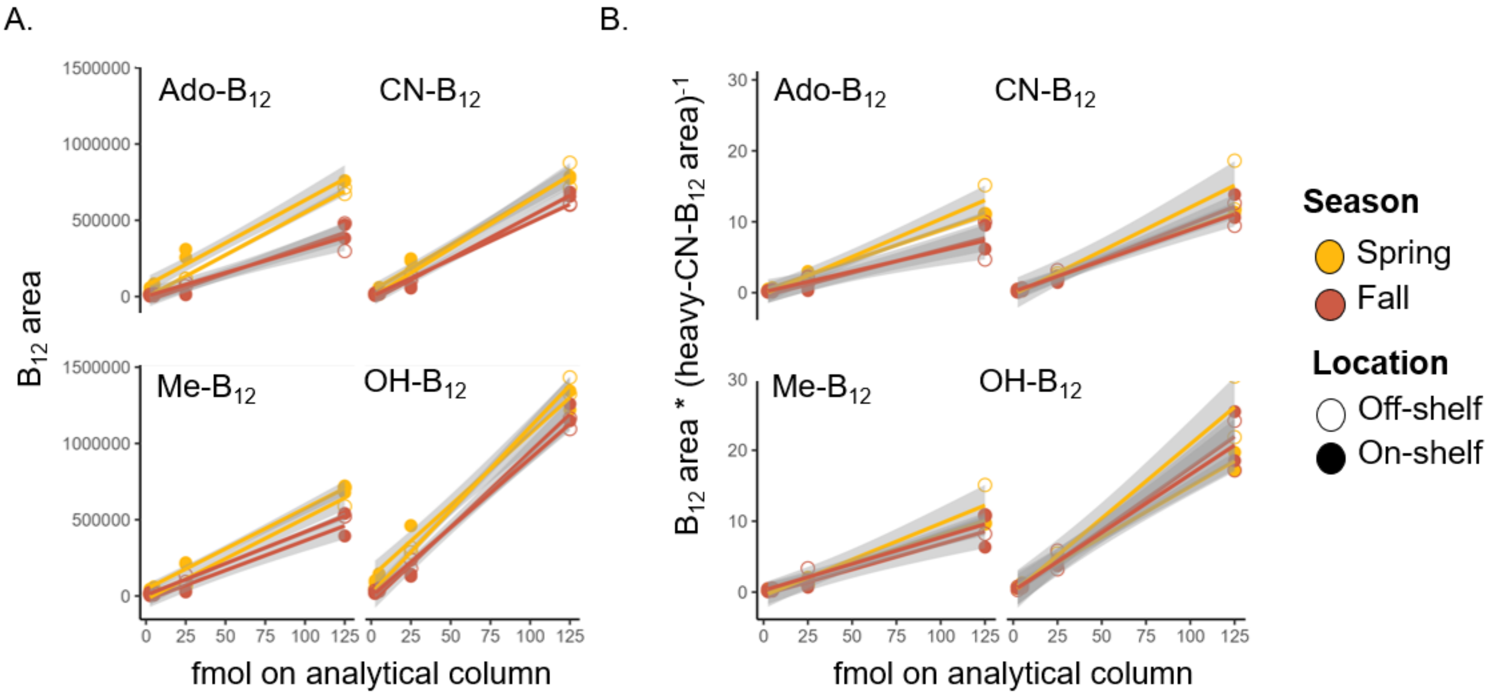

**Figure S3:** Nutrient additional bottle incubation (n= 3) collected on May 3rd, 2017, from HL02. (A) Chl a concentration (µg $L^{-1}$), and cells per mL of (B) >10 µm photosynthetic eukaryotic cells, (C) >3 µm but <10 µm photosynthetic eukaryotic cells, (D) <3 µm photosynthetic eukaryotic cells, (E) total photosynthetic eukaryotic cells, (F) *Synechococcus*, and (G) bacteria at $T_0$ and measured 4 days after addition of +$NO_3$ (10 µM), +$B_{12}$ (100 pM) and +$NO_3$+$B_{12}$ (10 µM, 100 pM). Different letters over data points indicate statistically significant differences (p-value < 0.05) between pairs based on the post-hoc Tukey's Test. Shared letters (i.e., a, b, c) indicate no significant difference, if no letters are displayed then the differences between treatments were insignificant.

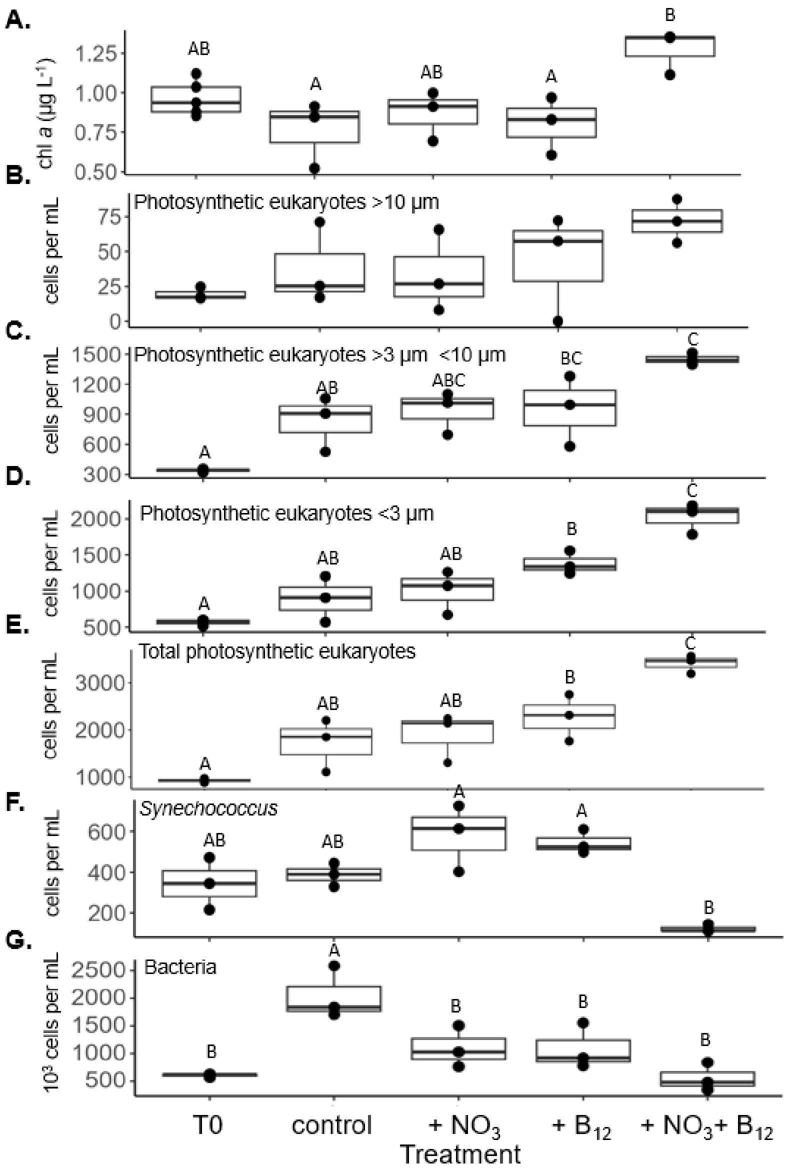

**Figure S4:** Concentrations (pM) of $B_2$ in (A) dissolved and (B) particulate phase over time in spring and fall.

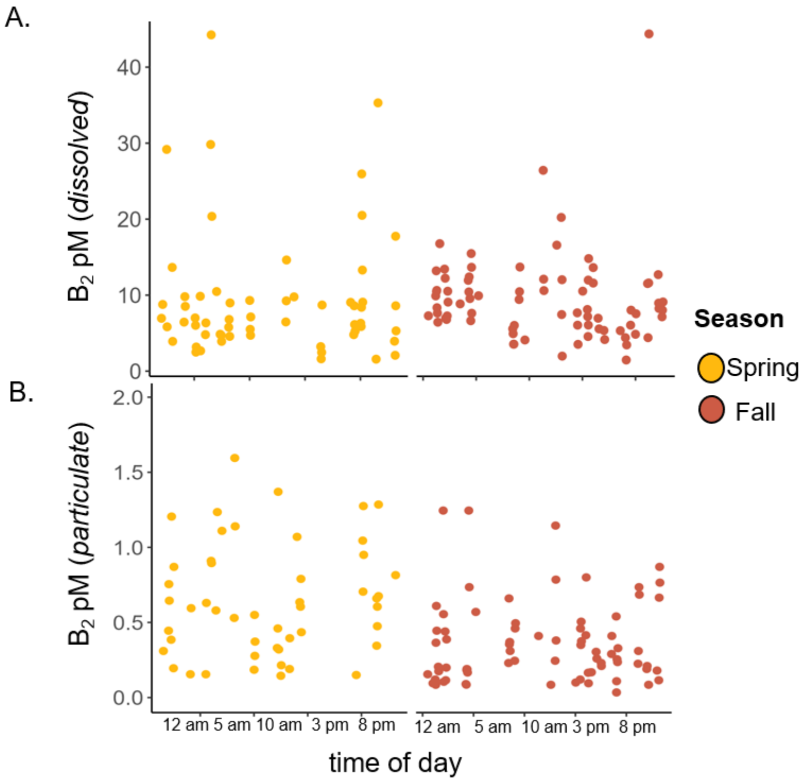
